# Contrasting patterns of microbial dominance in the *Arabidopsis thaliana* phyllosphere

**DOI:** 10.1101/2021.04.06.438366

**Authors:** Derek S. Lundberg, Roger de Pedro Jové, Pratchaya Pramoj Na Ayutthaya, Talia L. Karasov, Or Shalev, Karin Poersch, Wei Ding, Anita Bollmann-Giolai, Ilja Bezrukov, Detlef Weigel

**Affiliations:** Department of Molecular Biology, Max Planck Institute for Biology, 72076 Tübingen, Germany; Swedish University of Agricultural Sciences, 75007 Uppsala, Sweden; Centre for Research in Agricultural Genomics, 08193 Barcelona, Spain; ETH Zürich, 8093 Zürich, Switzerland; School of Biological Sciences, University of Utah, 84112 Salt Lake City, USA; University of Tübingen,Tübingen, Germany; University of Freiburg, 79085 Freiburg im Breisgau, Germany; University of Basel, 4001 Basel, Switzerland; University of Mainz, 55128 Mainz, Germany; Department of Evolutionary Biology and Environmental Studies, University of Zürich, 8057 Zürich, Switzerland

## Abstract

*Sphingomonas* is one of the most abundant bacterial genera in the phyllosphere of wild *Arabidopsis thaliana,* but relative to *Pseudomonas*, the ecology of *Sphingomonas* and its interaction with plants remains elusive. We analyzed the genomic features of over 400 *Sphingomonas* isolates collected from local *A. thaliana* populations, which revealed high intergenomic diversity, in contrast to genetically much more uniform *Pseudomonas* isolates found in the same host populations. Variation in *Sphingomonas* plasmid complements and additional genomic features suggest high adaptability of this genus, and the widespread presence of protein secretion systems hints at frequent biotic interactions. While some of the isolates showed plant-protective properties in lab tests, this was a rare trait. To begin to understand the extent of strain sharing across alternate hosts, we employed amplicon sequencing and a novel bulk-culturing metagenomics approach on both *A. thaliana* and neighboring plants. Our data reveal that *Sphingomonas* and *Pseudomonas* both thrive on other diverse plant hosts, but that *Sphingomonas* is a poor competitor in dying or dead leaves.

## Introduction

Most ecosystems, including host-associated microbiomes, are composed of a handful of common species and a wide assortment of rarer species (*1–3*). Common species are frequently implicated in direct interactions with the host and other microbes. For example in the human gut, the genera *Bacteroides* or *Prevotella*, which often occupy 20% or more of the entire community (*4*), modulate the host immune system and are particularly successful competitors of other microbes (*5*). Similarly, on and inside plant leaves, the Proteobacterial genera *Sphingomonas* and *Pseudomonas* are among the most common bacterial taxa, not only on the model plant *Arabidopsis thaliana* (*6–10*), but also on many other species across continents (*11–18*).

*Pseudomonas*-leaf interactions are widely-studied (*13, 19–21*), largely due to agricultural plant pathogens in the genus (*13*). *Sphingomonas*-leaf interactions, however, are not well-understood at a genetic level or population level, despite the ubiquity of *Sphingomonas* on plant leaves and increasing reports of plant beneficial members of this genus (*22–24*). *Sphingomonas* derives its name from its membrane-bound sphingolipids (*25*), structural and signaling molecules that are common in eukaryotes but found only in a few bacterial taxa (*4, 26*). These taxa include the previously mentioned *Bacteroides* and *Prevotella* of the gut, whose sphingolipids interact with the mammalian host and even influence host nutrition (*4, 27*). *Sphingomonas* also associate with plant roots (*23, 28, 29*) and seeds (*22*) and are common in soil and freshwater (*30*) among other habitats. Some strains can improve plant growth and abiotic stress tolerance in contaminated soils (*31*), and various others can promote plant growth through the production of growth regulators (*32*). *Sphingomonas* strains affect the abundances of other microbes (*6, 22, 33, 34*), and some protect against pathogenic bacteria (*22, 35*) or fungi (*23*).

In our previous studies of microbes colonizing local *A. thaliana* in southwest (SW) Germany, we also initially focused on characterizing *Pseudomonas* populations, and we use those results here as a benchmark for understanding *Sphingomonas*. We reported that *Pseudomonas* varied widely in bacterial load on individual plants, ranging from being nearly absent to very high titers (*8, 10, 19*). At the genomic level, 1,524 *Pseudomonas* isolates from local *A. thaliana* plants consisted primarily of a group of closely-related *P. viridiflava* (“OTU5” in the original publication and hereafter referred to as *Pv-ATUE5* (*36*)) that shared at least 99.9% nucleotide identity in their core genomes and the same diagnostic partial 16S rRNA gene sequence (*19*). Despite their similarity to each other, *Pv-ATUE5* strains differed widely in pathogenicity in a gnotobiotic system, and phylogenetic analysis suggested that subgroups of *Pv-ATUE5* diverged around 300,000 years ago, consistent with complex selective pressures that have not favored conquest by a single isolate (*19*).

In this work, we sought to characterize local *Sphingomonas* populations at the strain level to ask if, like *Pv-ATUE5*, a single lineage rules the local *A. thaliana* bacterial community, and to determine general genetic features of plant-associated *Sphingomonas*. We also extended our survey onto neighboring individuals of various plant species at the site and asked to what extent *Sphingomonas* and *Pseudomonas* strains common to *A. thaliana* are generalists, with selective pressures potentially shaped by life on multiple host plants. This work reinforces the notion that individual *Sphingmonas* strains likely have broad host ranges and that they have genetic features equipping them for a multitude of biotic interactions, suggesting major roles not only in the assembly of leaf communities but also in the general means by which plants sense and respond to non-pathogenic or beneficial microbes.

## Results

### Sphingomonas colonizes A. thaliana more consistently than Pseudomonas

A previous dataset of sequenced metagenomes for 176 *A. thaliana* rosette samples (*10*) allowed us to calculate the bacterial load for each rosette as the ratio of bacterial to plant sequence counts. We used these ratios to scale amplicon sequence variants (ASVs) of the V4 region of the 16S rRNA gene (hereafter rDNA) that had been prepared from the same DNA samples. This allowed us to quantify bacterial loads for each ASV, allowing us to overcome the compositionality problems inherent in relative abundance data (*37*). We first analyzed these ASVs at the genus level. *Sphingomonas* and *Pseudomonas* were approximately equal in abundance overall, and twice as abundant as the third most common genus (*Pedobacter*). However, *Pseudomonas* loads varied widely between samples, with a coefficient of variation (standard deviation / mean abundance) more than double that of *Sphingomonas*, which was among the most consistent genera (Fig. 1a). Impressively, the consistency of *Sphingomonas* colonization also extended to individual *Sphingomonas* ASVs that independently colonized most plants (Fig. 1b), suggesting that strains from each ASV may occupy a different niche on the leaf, or that different strains may reach an equilibrium as a community that limits growth. We named the most abundant *Sphingomonas* and *Pseudomonas* ASVs, for which the average abundance was several fold higher than the next most abundant ASV in each genus, SphASV1-V4 and PseASV1-V4, respectively.

**Fig. 1.**
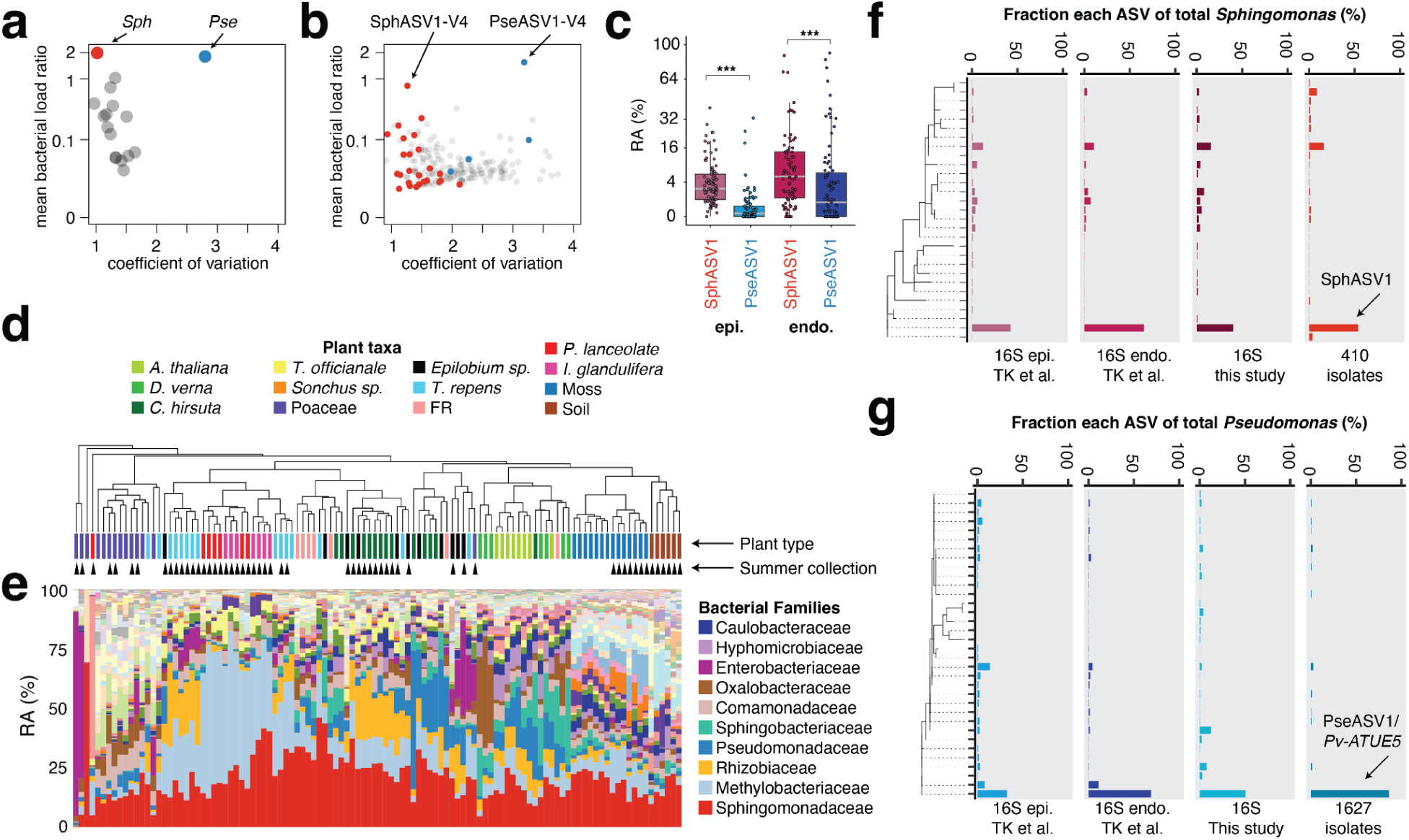
***Sphingomonas* and *Pseudomonas* are abundant and culturable, but differ in consistency of colonization. a**, *Sphingomonas* (red, *Sph*) has a similarly high bacterial load to *Pseudomonas* (blue, *Pse*) across 176 *A. thaliana* plants from (*10*), but among the lowest coefficients of variation (standard deviation / mean abundance) of other major genera (grey). **b**, Similar to **(a)**, but showing the V4 16S rDNA sequences that make up the most abundant genera. *Sphingomonas* ASVs (including SphASV1-V4) have on average low coefficients of variation, while *Pseudomonas* ASVs (including PseASV1-V4) have on average higher coefficients of variation. **c**, The median relative abundance (RA) of SphASV1 is higher than PseASV1 both epiphytically (epi.) and endophytically (endo.) (*** *p* < 0.001, FDR-adjusted Mann-Whitney U-test). **d)** Plant samples in the present study are clustered by the Bray-Curtis dissimilarity of the relative abundance of bacterial families. Summer collection indicated with black triangles. **e**, Stacked bars represent the relative abundances (RA) of bacterial families and correspond to the plant samples directly above in **(d)**. The colors identifying the top 10 bacterial families are shown in the legend. **f**, Relative abundances of *Sphingomonas* ASVs detected epiphytically (epi.) and endophytically (endo.) in TK et al. (*19*), this study, and in 410 locally-cultured isolates from multiple plant species. SphASV1 is indicated. **g**, Same as in **(f)**, but for *Pseudomonas*, with 1,627 isolates combining 1,524 isolates from (*19*) and 103 additional isolates from this study. PseASV1 is indicated.

### Dominant *Sphingomonas* and *Pseudomonas* ASVs are enriched endophytically

We reanalysed our previous dataset of slightly longer rDNA sequences in the V3V4 region (*19*), from samples in which we had separated *A. thaliana* leaf epiphytes from endophytes using surface sterilization that included a DNA-degrading bleaching step. We located longer ASVs containing an exact match to the SphASV1-V4 and PseASV1-V4 described above. We found for each a single match to a highly abundant V3V4 ASV, and we refer to the longer V3V4 rDNA sequences henceforth as SphASV1 and PseASV1. Unsurprisingly, PseASV1 uniquely matched the representative sequence previously used to define the abundant *Pv-ATUE5* lineage (*19*). SphASV1 was more abundant than PseASV1 in both the epiphytic and endophytic fractions of the leaf, and both PseASV1 and SphASV1were well-represented in the endophytic compartment (Fig. 1c), suggesting the capability to evade or tolerate the plant immune system and persist inside the leaf. SphASV1 was in fact the most abundant single ASV in the dataset. The combination of numerical dominance, endophytic enrichment, and high consistency of colonization drove our interests to characterize genetic and ecological features of this ASV and local *Sphingomonas* more generally.

### *Sphingomonas* has a wider host range than *Pseudomonas* in the phyllosphere

An abandoned train station in Eyach, Germany, supports a population of hundreds to thousands of *A. thaliana* plants that are predominantly winter annuals, and has been a source of material for several recent studies of *A. thaliana*-associated microbiomes (*6, 8, 19, 38*). We collected *A. thaliana* and the most common neighboring plants in spring, when most *A. thaliana* plants were green, mature, and flowering. We also collected common non-*A. thaliana* plants and bulk soil samples in late summer after all local *A. thaliana* vegetation had dried up and died, and the next cohort had not yet germinated. We reasoned that by late summer, microbes that had resided on spring *A. thaliana* must have long since migrated to survive on other plants or habitats (*13, 39, 40*) or died along with their host, and thus at this later time point, isolates collected from other plant species or soil could not be recent migrants from *A. thaliana* .

We chose each non-*A. thaliana* plant species based solely on its abundance at the site and not by any prior preference for certain plant species, and we therefore chose some plants that we were unable to name, despite being able to confidently classify and recognize different individuals by their morphology. For each plant species, we pooled at least one entire leaf from at least 6 independent individuals per sample, collecting 7 independent samples per species. Although this pooling strategy that spanned many plants per sample precluded analysis of variation between individual plants, it maximized our ability to broadly survey the site (Extended Data Fig. 1). After bringing leaf samples back to the lab, we surface-sanitized them in 75% ethanol for 45-60 seconds (*41*) and macerated them in phosphate buffered saline (PBS) buffer with a mortar and pestle. Soil samples were directly mixed with PBS buffer. We then mixed the fresh lysates with glycerol to make -80°C freezer stocks that cryo-protected the live bacteria (Methods).

We first extracted metagenomic DNA directly from aliquots of the frozen lysates using a custom bead-beating based protocol as previously described (*10, 19*), and prepared 16S rDNA amplicon libraries spanning the V3 and V4 regions (Methods) using peptide nucleic acids (PNAs) to reduce organelle amplification (*42*). The residual chloroplast sequences in each sample were consistent within each plant pool, reaffirming that we had not mistakenly pooled leaves from different plant taxa (Extended Data Fig. 2). Unfortunately, we could not obtain sufficient bacterial reads from dandelion and thistle due to a natural mutation in their chloroplast sequences that made our PNAs ineffective (*43*), leading to an overabundance of chloroplast sequences. To contrast bacterial communities across the sampled plant hosts, we binned the ASVs into bacterial families, and clustered samples by their pairwise Bray-Curtis dissimilarity (Fig. 1d). This revealed substantial similarities in bacterial family membership between groups of samples, with clear plant clustering both by sampling season and also by some plant taxa, consistent with previous publications linking plant genotype and seasonal effects to bacterial community composition (*14, 44*). The family Pseudomonadaceae, of which 92.9% belonged to the genus *Pseudomonas*, reached high relative abundances in the *A. thaliana* phyllosphere, but also in some other hosts—particularly the other Brassicaceae *Draba verna* and *Cardamine hirsuta* (Fig. 1e). Sphingomonadaceae, of which 80.7% corresponded to the genus *Sphingomonas*, was ubiquitous and abundant across all plant hosts (Fig. 1e, Extended Data Fig. 3). Across all plants, PseASV1 and SphASV1 accounted for a substantial fraction of the reads from *Pseudomonas* and *Sphingomonas* (44.7% and 39.4%, respectively) (Fig. 1f, 1g).

### Cultured bacterial populations resemble those on wild leaves

*Pseudomonas* isolates from local *A. thaliana* populations in Germany, the majority of which have the PseASV1 16S rDNA sequence, have been previously characterized (*19*). To investigate *Sphingomonas* genome diversity across host species, and likewise to characterize the genomic features associated with highly abundant *Sphingomonas* groups, we cultured *Sphingomonas* from frozen plant lysates using both remaining *A. thaliana* lysates from (*19*) as well as from *A. thaliana* plus diverse plant hosts in the present study. We enriched for *Sphingomonas* using rich LB media supplemented with cycloheximide and 100 µg/mL streptomycin, an antibiotic to which most *Sphingomonas* is resistant due to a natural mutation in the *rpsL* gene (*45*), and isolated 410 *Sphingomonas* colonies. Using LB supplemented with cycloheximide and 100 µg/mL nitrofurantoin as previously described (*19*), we also isolated an additional 103 Pseudomonads. We generated draft genome assemblies with an average of 65-fold genome coverage and BUSCO (*46*) completeness of at least 85% (Methods). We annotated open reading frames using Prokka (*47*) and extracted 16S rDNA sequences using barrnap (*48*). We first analyzed the V3 and V4 regions of the 16S rDNA, which allowed us to match the genomes to our existing culture-independent ASVs. Critically, for both *Pseudomonas* and *Sphingomonas*, we recovered isolates in relative abundances consistent with those from culture-independent surveys (Fig. 1f, 1g), demonstrating that culturing did not introduce substantial biases in recovery rates, and that we had sequenced a sufficient number of isolates to capture broad patterns of diversity on leaves.

### *Sphingomonas* 16S rDNA sequence similarity belies high genomic diversity

Previously we observed low genomic diversity among PseASV1/*Pv-ATUE5* strains isolated from *A. thaliana* (*19*). To similarly evaluate genomic diversity for an analogous set of *Sphingomonas*, we selected all the *Sphingomonas* isolates in our collection that both came from *A. thaliana* and had the SphASV1 16S rDNA sequence (representing 174 with SphASV1 out of 340 total *A. thaliana* isolates). We compared the genomes to each other using MASH (*49*), which decomposes genomes into *k*-mers and calculates a distance based on the fraction of shared *k*-mers. As a comparator, we also included a representative set of 99 diverse PseASV1-associated genomes isolated from *A. thaliana*, comprising 82 strains from (*19*) that were at least 0.1% different in MASH distance from all others across the core genome, and an additional group of 17 PseASV1-associated strains from this study. We converted the MASH distances to a similarity score between 0 (least similar) and 100 (identical) which closely corresponds to average nucleotide identity (ANI) (Methods) (*50, 51*). When PseASV1-associated genomes were compared to each other, all pairs had MASH similarities > 96. However, pairwise comparisons between SphASV1-associated *Sphingomonas* genomes had MASH similarities as low as 81 and averaging 89, indicating that the V3V4 region of the 16S rDNA sequence was a relatively poor predictor of genome similarity (Fig. 2a, 2b). Generally, the full 16S rDNA sequence (as opposed to one or two variable regions) provides increased discriminatory power between strains (*52*). To see the extent of this, we therefore clustered the full-length 16S rDNA sequences extracted from the genomes into operational taxonomic units (OTUs) at 99.5% identity. This yielded no additional subgroups for PseASV1-associated strains, but partitioned SphASV1-associated strains into 6 subgroups, five of which included more than 1 strain and could be compared (Fig. 2a). Genomes within these five subgroups were more similar, with average intragroup MASH similarities of 90, 91, 91, 96, and 97. All of these values were lower than for *Pseudomonas*, further confirming that 16S rDNA sequences were less informative for local *Sphingomonas* than for local *Pseudomonas* (Fig. 2b). We further explored the ability of the gyrase B (*gyrB*) gene, a commonly-used high-resolution phylogenetic marker, to distinguish strains, and extracted *gyrB* sequences from the assembled genomes (Fig. 2a). *gyrB* outperformed 16S rDNA as a strain-specific marker, and each *Sphingomonas* shared its *gyrB* sequence with on average 1.6 other SphASV1-associated strains, while each *Pseudomonas* shared its *gyrB* sequence with on average 4.3 other PseASV1-associated strains.

**Fig. 2.**
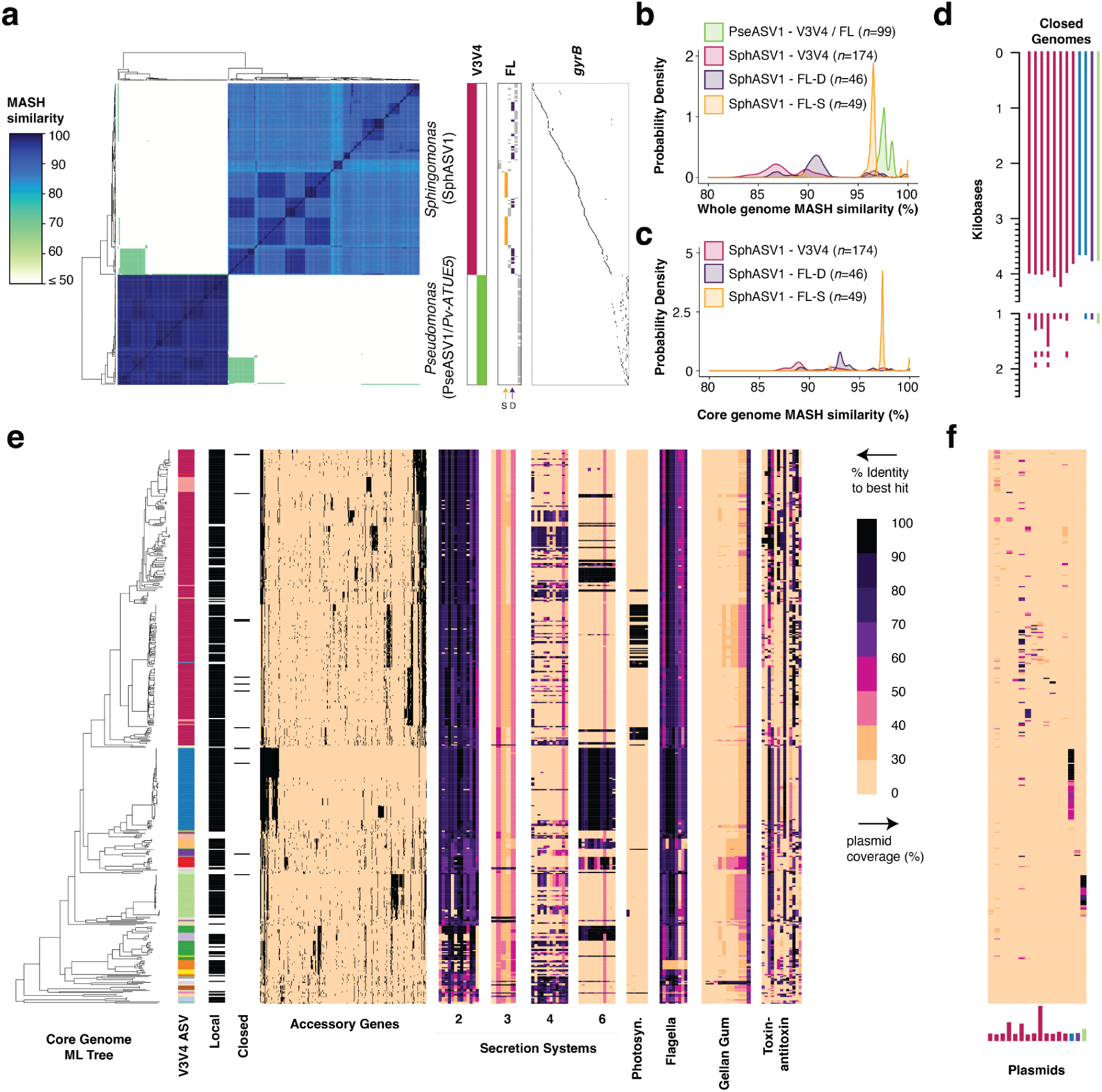
***Sphingomonas* 16S rDNA sequences belie high genomic diversity. a**, Heatmap of MASH similarity between 99 *Pseudomonas* PseASV1/*Pv-ATUE5* genomes and 174 *Sphingomonas* SphASV1 genomes. The heatmap aligns to the vertical bars to the right, which indicate which genomes contain the V3V4 16S rDNA ASV used to define PseASV1 and SphASV1 (‘V3V4’, magenta and green), which genomes have different full length 16S rDNA OTU (‘FL’), and which genomes differ in their *gyrB* fragments (‘gyrB’). A diverse group (‘D’) and a similar group (‘S’) of genomes sharing a full length OTU sequence are indicated by arrows underneath the ‘FL’ panel. **b**, Density plot showing the distribution of MASH similarities from the heatmap in **(a)**, subset by bacterial genus and 16S rDNA grouping. V3V4, variable regions 3 and 4; FL, a full length OTU; FL-D and FL-S, the groups of highly diverse and highly similar genomes, respectively, sharing a full length OTU sequence as indicated in **(a)**. **c**, similar to **(b)** but showing MASH similarities calculated only on an alignment of 274 core genes. **d**, Base map of the 12 closed genomes, colored by their V3V4 16S group, showing plasmids to scale. **e**, Core genome maximum likelihood (ML) tree of 410 *Sphingomonas* isolates from this study and 70 from RefSeq based on 274 core genes, with genomic features indicated by aligned heatmaps. ‘V3V4’, color map of the different V3V4 ASV sequences associated with each genome (white if unknown); ‘FL OTU’, color map of the 99.5% identity OTUs associated with SphASV1 and and closely-related ASVs; ‘Local’, 410 genomes sequenced and isolated in this study are indicated by black; ‘Accessory Genes’, presence/absence matrix of all genes not used to compute the ML tree; ‘Secretion Systems’, BLAST results of each genome against genes of the type 2, 3, 4, and 6 secretion systems, with percent identities as indicated by the color legend. ‘Photosyn’, genes involved in anoxygenic photosynthesis; ‘Flagella’, genes involved in building an operating flagellum; ‘Toxin-antitoxin’, genes associated with toxin-antitoxin pairs. **f**, ‘Plasmids’, percent coverage of plasmid sequences from the closed genomes, with the relative size and ASV background of each plasmid indicated by the length and color of the bars.

We initially suspected that the relatively high overall genome diversity of SphASV1-associated isolates compared to PseASV1-associated isolates might be due to high variation in the accessory genomes—specifically the differential presence of plasmids. We noted that some publicly-available complete genomes of *Sphingomonas* contained several plasmids, and after closing 12 of our genomes with long-read sequencing (Methods), we detected a total of 16 plasmids with up to three per genome and comprising up to 14% of the total genome size (Fig. 2d). To see if plasmids and mobile elements were driving low *Sphingomonas* genome similarity, we considered only conserved sequences in genomes by first creating a “soft” core genome using PanX (*53*) to identify open reading frames present in at least 70% of genomes, resulting in a set of 274 shared genes. MASH comparisons of core genomes of SphASV1-associated isolates (Fig. 2c) were still more diverse than even whole genomes of PseASV1-associated isolates. Thus, despite the pervasiveness of the SphASV1 sequence in our dataset, one cannot reliably extrapolate what this ASV sequence means at the level of genomic content.

### *Sphingomonas* genomes reveal adaptations for competitive life in the phyllosphere

To explore relatedness of SphASV1-associated strains, and how they compare with other *Sphingomonas*, we calculated maximum-likelihood (ML) “soft” core genome phylogenies (*53*) from all 410 local *Sphingomonas* isolates (340 from *A. thaliana* and 70 from other local plant hosts), along with 70 sequence-related isolates from NCBI RefSeq (*54*). We intentionally did not include highly divergent RefSeq isolates, many of which were isolated from non-plant environments. Gene presence or absence in the accessory genome (Fig. 2e) tracked well with differences in the core genome; this was tested by correlating pairwise Euclidean distances calculated on the presence/absence matrix of accessory genes to MASH distances calculated on core genes (R^2^=0.74). We compiled a short list of *Sphingomonas* genes that had a high likelihood, based on literature, to improve survival among competing bacteria, or to facilitate interaction with a plant host, and made a custom database of *Sphingomonas* protein sequences from RefSeq. We searched for these features in our genomes by aligning the assemblies using BLASTX (*55*), considering alignments to be matches if they had at least 30% amino acid identity over at least 60% of the protein length. While nearly all isolates had diagnostic genes for the type 2 secretion system, and genes for the type 4 and type 6 secretion systems were common among some clades including those of SphASVI, only a handful of less abundant strains seemed to potentially have a type 3 secretion system (Fig. 2e). Flagellar motility is common. A group of SphASV1 genomes has a full suite of genes for anoxygenic photosynthesis, a fascinating feature that can supplement heterotrophic energy production and likely improves survival in well-illuminated and nutrient-poor conditions such as the phyllosphere (*56–59*). All genomes were rife with toxin-antitoxin systems, likely to stabilize plasmids or superintegrons (*60*). Indeed, we found widespread evidence of plasmids in our draft genomes by using minimap2 (*61*) to search for alignments to the 16 circularized plasmid sequences from our 12 complete genomes (Fig. 2d). Considering alignments covering at least 30% of the length of a plasmid as a positive hit, 250 (60%) of our isolates showed signatures of one or more of these 16 plasmids (Fig. 2f).

To increase confidence in the preceding gene searches in our draft genomes, we repeated the analysis comparing the 12 closed genomes to their draft genome counterparts. Both draft and complete versions of each genome clustered tightly in a ML tree, and showed essentially the same presence/absence patterns (Extended Data Fig. 4), demonstrating that draft genomes were sufficient for analysis of gene content at this level of detail.

### Some Sphingomonas attenuate Pseudomonas virulence in A. thaliana

Besides reaching similarly high abundances in the same leaves (Fig. 1a-b), both *Sphingomonas* and *Pseudomonas* grow on many similar substrates *in vitro* (*35*), suggesting potential microniche overlap in the phyllosphere. Previous work revealed that certain strains of *Sphingomonas*, in particular *S. melonis* Fr1, can ameliorate symptoms in *A. thaliana* leaves caused by pathogenic *Pseudomonas* and *Xanthomonas* in a gnotobiotic system (*24, 35, 62*). More recently, a strain of *S. melonis* was demonstrated to protect rice against the bacterial seedling blight pathogen *Burkholderia plantarii* (*22*). We first sought to screen some of our local isolates for potential plant-protective activity against virulent *Pv-ATUE5* strains in a gnotobiotic 24-well plate agar system as we used previously (*19*).

We germinated an *A. thaliana* accession endemic to our field site, Ey15-2, on half-strength MS solid media in the presence of each of 19 strains of local *Sphingomonas* that represented much of the diversity in our collection, as well as *S. melonis* Fr1. Following 10 days of co-cultivation, we challenged the plants with *Pseudomonas*, either the virulent local *P. viridiflava* strain *Pv-ATUE5*:p25c2 (*19*) or the model pathogen *P. syringae* (*Pst*) DC3000, and monitored plant health over the next 6 days by measuring green pixels (Methods, Extended Data Table 3). *Pst* DC3000 slowed plant growth compared to buffer control, while *Pv-ATUE5*:p25c2 killed plant tissues (Fig. 3a-d, Extended Data Fig. 5). Surprisingly, seedlings germinated in the presence of *S. melonis* Fr1 in these conditions were consistently stunted compared to those germinated in buffer or those germinated with any other *Sphingomonas* (Fig. 3a, 3c, Extended Data Fig. 5). However, despite this negative effect on growth, *S. melonis* Fr1 protected plants from the worst effects of *Pv-ATUE5*:p25c2, with infected plants not dying but instead growing more slowly, not significantly different from plants treated with the less deadly *Pst* DC3000 across two replicates (FDR-adjusted Mann-Whitney U-test, *p* > 0.05). Our local *Sphingomonas* strain *SphATUE*:S139H133 also protected plants, reducing *Pv-ATUE5*:p25c2 virulence such that it was no worse than *Pst* DC3000 across two independent experiments (Fig. 3a-b, Extended Data Fig. 5) (FDR-adjusted Mann-Whitney U-test, *p* > 0.05). Unlike *S. melonis* Fr1, *SphATUE*:S139H133 did not stunt growth.

**Fig. 3.**
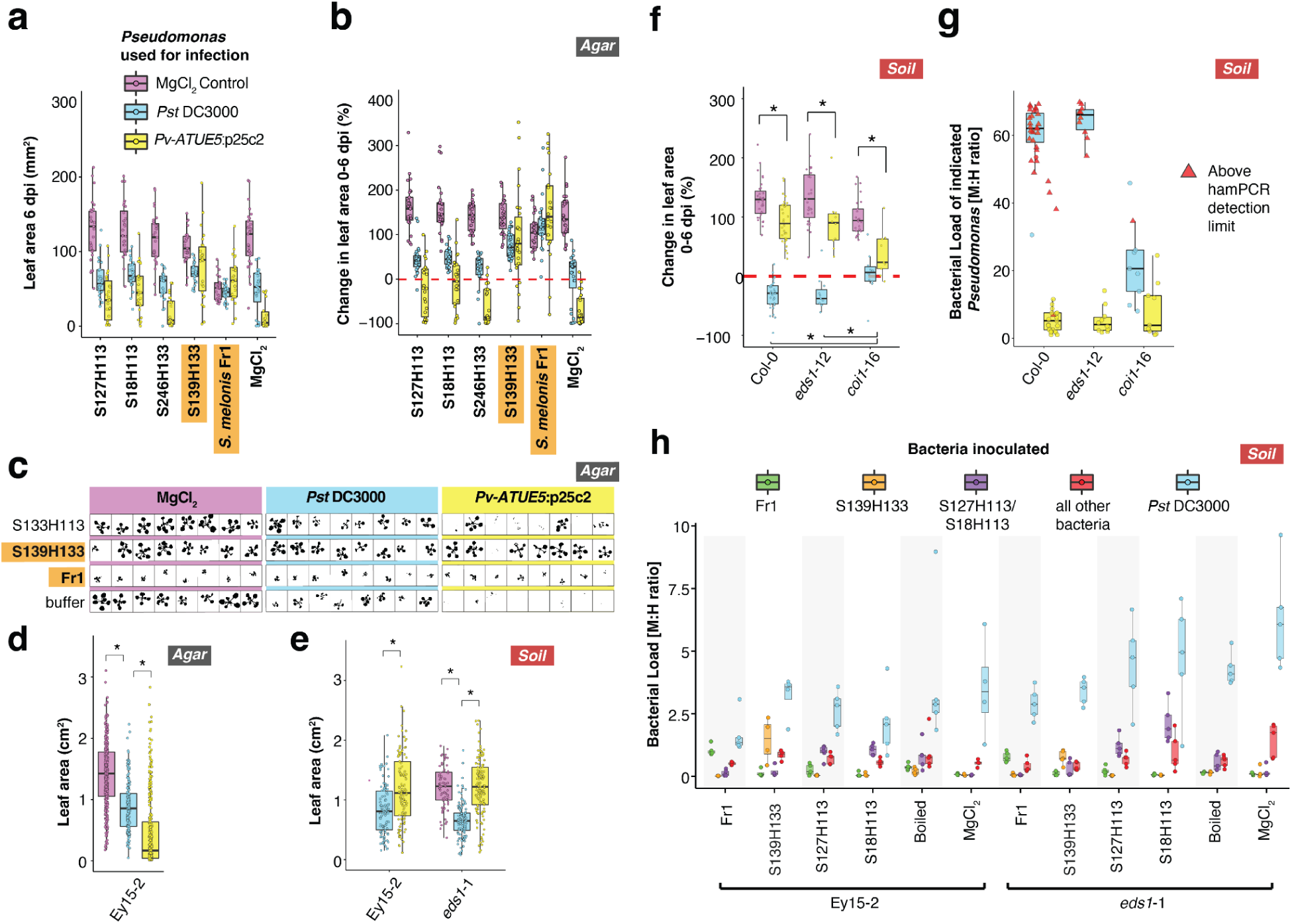
**Some *Sphingomonas* offer protection against *Pseudomonas*. a-d**, Local *Sphingomonas* isolates, *S. melonis* Fr1 (Fr1), or MgCl_2_ buffer were used to pre-treat *A. thaliana* Ey15-2 seeds germinating in 24-well agar plates. On day 10, seedlings were challenged with a MgCl_2_ buffer control, *Pst* DC3000, or *Pv-ATUE5:p25c2* and monitored for 6 days after infection (*n* ≥ 6 per condition). Orange-highlighted *Sphingomonas* insignificant difference between infected and buffer-treated plants (Mann-Whitney U-test with FDR adjustment, *p* > 0.05). **a**, Absolute size of rosettes at 6 dpi. **b**, Percent change in rosette size between 0-6 dpi. **c**, Representative silhouettes at 6 dpi of seedlings grown in 24-well plates. **d**, Rosette size by infection treatment, regardless of *Sphingomonas* pretreatment. * = P < 0.001 (Mann-Whitney U-test with FDR adjustment). **e-g**, Three local *Sphingomonas* isolates, *S. melonis* Fr1, boiled *Sphingomonas*, or buffer were sprayed on 2-week old seedlings of Ey15-2 or *eds1*-1, and 4 days later, live *Pseudomonas* or MgCl_2_ buffer was inoculated. **e**, Absolute sizes of rosettes for different *Pseudomonas* treatments, regardless of *Sphingomonas* treatment. * = *p* < 0.001 (Mann-Whitney U-test with FDR adjustment). **f**, Percent change in rosette size between 0-6 dpi for three genotypes in the Col-0 background infected with Pseudomonas or buffer control and grown on soil. **g**, Bacterial load ratio in plants infected with *Pst* DC3000 as determined by hamPCR. Note that the color legend represents bacteria quantified, not the bacteria used for treatment. **h**, Bacterial load ratio in plants infected with *Pst* DC3000 as determined by hamPCR. Note that the color legend represents bacteria quantified, not the bacteria used for treatment.

We also observed the stunting phenotype of *S. melonis* Fr1 in the Col-0 accession (not shown) for which a protective effect of *S. melonis* Fr1 had been previously reported (*62*). Although the authors reported that the plant transcriptomic response to protective *S. melonis* Fr1 involved induction of a set of defense genes overlapping those induced by *Pst* DC3000 (*62*), they did not observe any negative effects on plant growth (*24, 62*), and we therefore suspect that the unexpected plant stunting caused by *S. melonis* Fr1 in our experiments might be due to differences in our gnotobiotic system or in the growth media.

To test whether protective effects might be less apparent in the agar system, we directly tested plants grown in more natural conditions on soil. We therefore set up a similar experiment in which 2-week old Ey15-2 seedlings were grown on potting soil and sprayed first with one of four *Sphingomonas* strains, including the protective *SphATUE*:S139H133 and *S. melonis* Fr1, and a boiled *Sphingomonas* control. Four days later we challenged the plants with *Pseudomonas* sprayed at high concentrations (O.D._600_ = 1.0), using experimental conditions known to produce robust symptoms (*63*). We also included *enhanced disease susceptibility 1* (*eds1*-1, Ws-0 background) as an infection control because this mutant, defective for numerous defense responses mediated by salicylic acid (SA), is hypersusceptible to *Pst* DC3000 (*64*). At 5 days post infection (dpi), we observed classic *Pst* DC3000 symptoms on most plants including chlorotic leaves, stunted growth, and increased anthocyanin at the apical meristem (*63*), especially on *eds1*-1 plants. However *Pv-ATUE5*:p25c2, which was consistently deadly on agar plates, did not produce any obvious symptoms on soil-grown *A. thaliana* and did not greatly affect rosette size of even *eds1*-1 mutants during the time span of the experiment (Fig. 3e).

This unexpectedly weak *Pv-ATUE5*:p25c2 virulence prompted us, before proceeding with further analyses, to additionally test the *A. thaliana* mutant *coronatine-insensitive 1* (*coi1-*16), which is more susceptible to *P. viridiflava* (*65*) due to defects in defense responses mediated by jasmonic acid (JA). We therefore infected both *Pv-ATUE5*:p25c2 and *Pst* DC3000 on *coi1*-16 (Col-0 background), *eds1*-12 (Col-0 background) (*66*), and wild-type Col-0 using the same strong infection conditions.

We again observed no obvious discoloration or disease symptoms on any plant infected with *Pv-ATUE5*:p25c2, including the *coi1*-16 mutant, after 5 days. However, in this experiment *Pv-ATUE5*:p25c2 did significantly retard the growth of rosettes compared to a 10 mM MgCl_2_ control in all genetic backgrounds (Mann-Whitney U-tests with FDR adjustment, p < 0.01), in agreement with reports of *P. viridiflava* virulence on soil (*67, 68*), and the magnitude of this effect was strongest in the *coi1*-16 mutant (Fig. 3f), consistent with JA increasing resistance to *P. viridiflava* (*65*). In contrast, *Pst* DC3000 induced strong chlorosis on both Col-0 and *eds1*-1, but not on *coi1*-16, and retarded the growth of all plants, with *coi1*-16 being the least affected (Fig. 3f), consistent with the *coi1*-16 mutant being more resistant to *Pst* DC3000 (*69*). In addition to scoring disease symptoms and plant growth, we estimated *Pseudomonas* load in the mutants using hamPCR (*70*), a quantitative amplicon sequencing approach that derives bacterial load by relating abundance of 16S rDNA to a single-copy host gene. For Col-0 and *eds1*-1 plants, the load of *Pst* DC3000 exceeded the limits of robust quantification, while it was much lower on resistant *coi1*-16 plants, as expected. *Pv-ATUE5*:p25c2 loads were far lower than *Pst* DC3000 on all plants, explaining the milder virulence phenotype.

In the protection experiment on Ey15-2 and *eds1*-12 plants, we did not observe a consistent protective effect of any local *Sphingomonas* strain against *Pst* DC3000 symptoms on soil. However, for both Ey15-2 and *eds1-*1 plants, pre-treatment with *S. melonis* Fr1 did result in larger plants than pre-treatment with boiled *Sphingomonas* or with buffer (FDR-adjusted Mann-Whitney U-test, *p* < 0.05, Extended Data Fig. 6). In contrast to the agar system, *S. melonis* Fr1-treated Ey15-2 plants grown on soil were not stunted (Extended Data Fig. 6). To confirm that *Sphingomonas* was still present at the end of the experiment, and if pathogen titers had been affected by *Sphingomonas*, we quantified bacterial communities in the leaves of 8 plants per condition at 5 dpi using hamPCR (*70*). We observed the inoculated *Sphingomonas* 16S rDNA sequences enriched in each endpoint sample, as expected (Fig. 3g). Samples pre-treated with *S. melonis* Fr1 supported less *Pst* DC3000 proliferation than those pre-treated with other *Sphingomonas* of buffer in both the Ey15-2 and *eds1*-1 backgrounds, although the difference was only significant in *eds1*-1 (Mann-Whitney U-test, p<0.05). However, we found no evidence that other isolates, including the local Sph139H133, which was protective on agar, reduced pathogen titers in plants grown on soil in these strong infection conditions.

### *Sphingomonas* and *Pseudomonas* strains thrive on multiple plant species

*Sphingomonas* and *Pseudomonas* are generalists, with no known exclusive hosts of any given strain (*13*). As a first step towards determining the breadth of hosts in our study area, we compared *Sphingomonas* and *Pseudomonas* ASVs across local plant species. While SphASV1 was more abundant overall in the spring collection, it was consistently detectable on most plant species in both seasons. Plant taxa colonized by SphASV1 were also frequently colonized to appreciable levels by multiple other *Sphingomonas* ASVs (Fig. 4a). In contrast, PseASV1 was relatively more abundant on *A. thaliana, C. hirsuta* and *D. verna* - all from the family Brassicaceae - and less abundant, though still easily detectable, on other taxa (Fig. 4b). PseASV1 was especially enriched in the spring, which matches our previous finding of more *Pv-ATUE5* isolates from spring vs. winter collected *A. thaliana* (*19*).

**Fig. 4.**
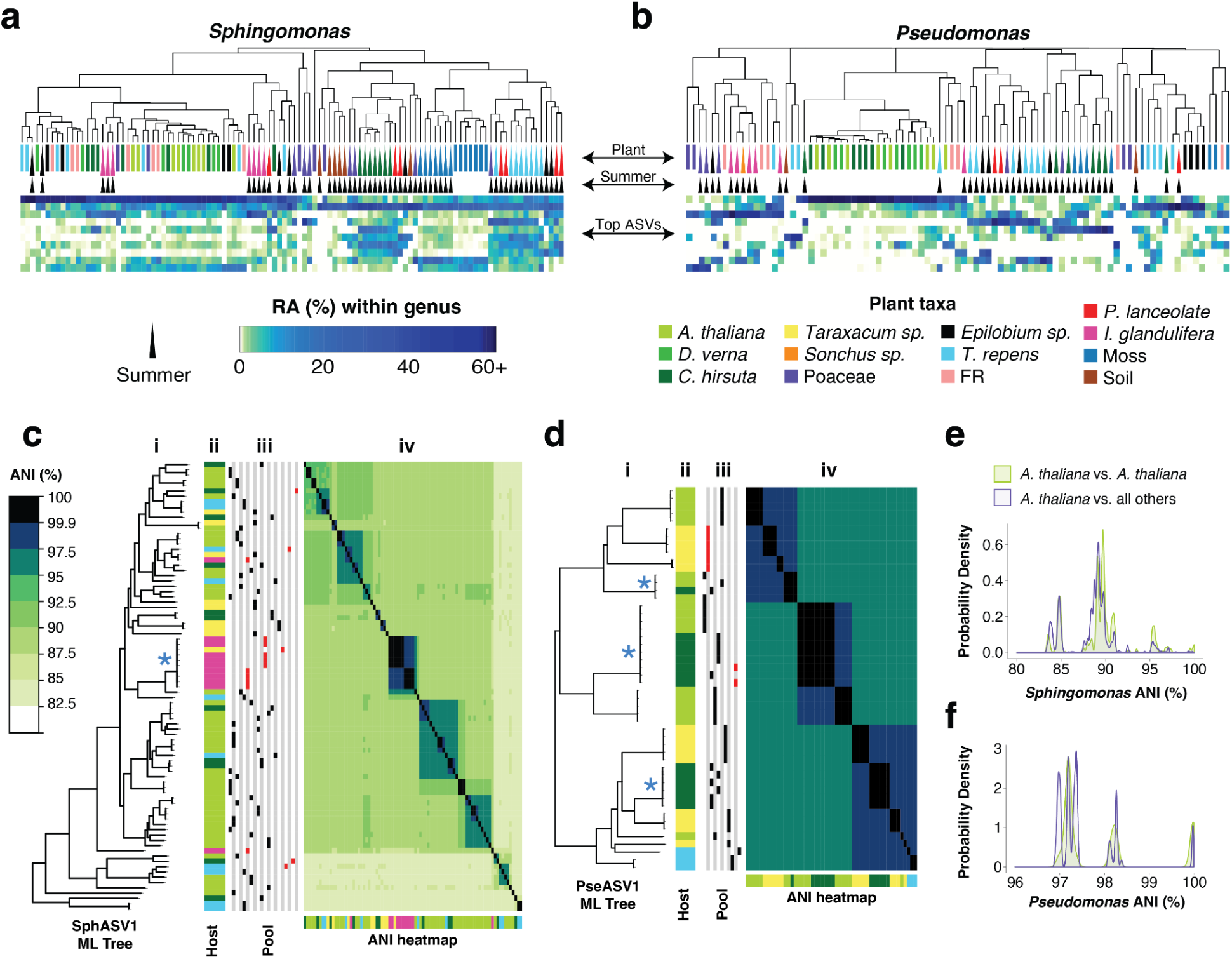
**Isolates from other plant species match those isolated from *A. thaliana*. a-b**, Host species (top) from 2018 clustered by Bray-Curtis dissimilarities between relative abundances (RA) for the 10 most abundant 16S rDNA ASVs for each genus. The legend below the Fig.s is shared for both panels. **a**, *Sphingomonas* ASVs, with SphASV1 as the top row in the heatmap. **b**, Same as **(a)**, but for *Pseudomonas* ASVs. **c**, Heatmap revealing genome ANI of all pairwise comparisons between SphASV1 isolates. **i:** Isolates arranged by a maximum likelihood (ML) core genome tree. The asterisk approximately in the middle of the tree represents the position of a group of isolates sharing > 99.9% ANI that derives from different sample pools, as indicated in columns **(iii)** to the right. **ii**: plant host of origin is indicated by color, corresponding to the plant legend in **(a-b**). **iii**: Isolates from the same sample pool are indicated by red (summer) or black (spring) marks in the same vertical column. **iv**: heatmap showing pairwise ANI values. **d**, Same as **(c)**, but for PseASV1. **e**, Distribution of pairwise intra-host isolate ANI compared to inter-host isolate ANI for SphASV1. **f**, Same as **(e)**, but for PseASV1 isolates.

Because a single ASV can comprise many different strains, the presence of the same ASV does not necessarily mean that the same specific strains are present on multiple host plants. To test for strain sharing across hosts, we compared PseASV1 and SphASV1 genomes of isolates from *A. thaliana, T. officinale*, *T. repens*, *C. hirsuta*, and *I. glandulifera* (Fig. 4c, 4d). These species were chosen because we had been able to easily isolate both *Pseudomonas and Sphingomonas* from the same plant pool lysates prepared for each of these species. We compared the genomes’ average nucleotide identity (ANI) because this commonly-used metric is robust to the incompleteness of draft genomes (*50*) and is a standard metric for defining bacteria relatedness and defining species (*51*). While 29 out of the 86 SphASV1 isolates from the 2018 harvest shared at least 99.9% ANI with at least one other isolate, with only one exception these highest-similarity isolates came from the same pool of plants, implying that there was little evidence for the same clonal *Sphingomonas* lineage independently colonizing different plant individuals. To exclude potential clones from the same plant, we recalculated all pairwise genome similarities between SphASV1 isolates from different lysates. We compared similarity within *A. thaliana* isolates (intra-host isolate similarity) and between *A. thaliana* isolates and those from any other plant species (inter-host isolate similarity). A higher intra-host similarity would be evidence of host-specificity, possibly due to unique selective forces within *A. thaliana*, or easier migration between individuals. We found that the distribution of intra-host ANI values closely matched that of inter-host ANI values (Fig. 4e), with the intra-host ANI being only marginally higher (0.9%, Mann-Whitney U test, *p* < 0.001).

PseASV1 genomes were more similar to each other than SphASV1 genomes (Fig. 2a). Every isolate shared at least 97.5% ANI with at least one isolate from a different plant species, and 47 of the 50 isolates shared at least 99.9% ANI with at least one other isolate (Fig. 4d), although as with SphASV1, these highest-similarity isolates tended to come from the same plant pool. However, three groups of highest-similarity isolates were shared across different plant pools, different species, and even different seasons (Fig. 4d). This is consistent with previous observations that *Pv-ATUE5* strains are common and persistent on diverse *A. thaliana* populations (*19*), and demonstrates that other local plant species host these strains as well. As with SphASV1 isolates, the intra-host ANI values followed a similar distribution to inter-host ANI values, with intra-host isolate ANI values again being marginally higher (0.4%, Mann-Whitney U-test, *p* < 0.001, Fig. 4f).

### Bulk culture metagenomics reveals strain sharing across plant species

From our genome-sequenced isolates, we had found that closely related *Sphingomonas* and *Pseudomonas* strains could colonize diverse hosts. To broaden our survey and extend these observations, we adopted a time and cost-efficient approach to enrich each genus in bulk from plant lysates and sequence the enriched pool as a metagenome. We plated 50 μL of glycerol stock from each lysate, corresponding to ∼5 mg of original plant material, on either selective *Pseudomonas* or *Sphingomonas* medium and grew colonies *en masse* (Fig. 5a, Extended Data Fig. 7). This quantity of lysate was chosen empirically for our samples to avoid colonies merging into competitive lawns and to allow observation of discrete colonies. As a control for lysate viability and a rough reference of bacterial load, we also cultured bacteria from lysates on non-selective rich LB medium. We counted the colonies on each plate after 2 days at room temperature for *Pseudomonas* or mixed bacteria on LB, or after 7 days for slower-growing *Sphingomonas* (Fig. 5b)(Methods). As expected, rich LB medium yielded more colonies than the two selective media, although counts on LB likely greatly underestimate the total bacteria population in each sample, not only because a single rich medium does not meet nutritional demands for all bacterial taxa, but also because fast-growing colonies may overgrow or suppress others.

**Fig. 5.**
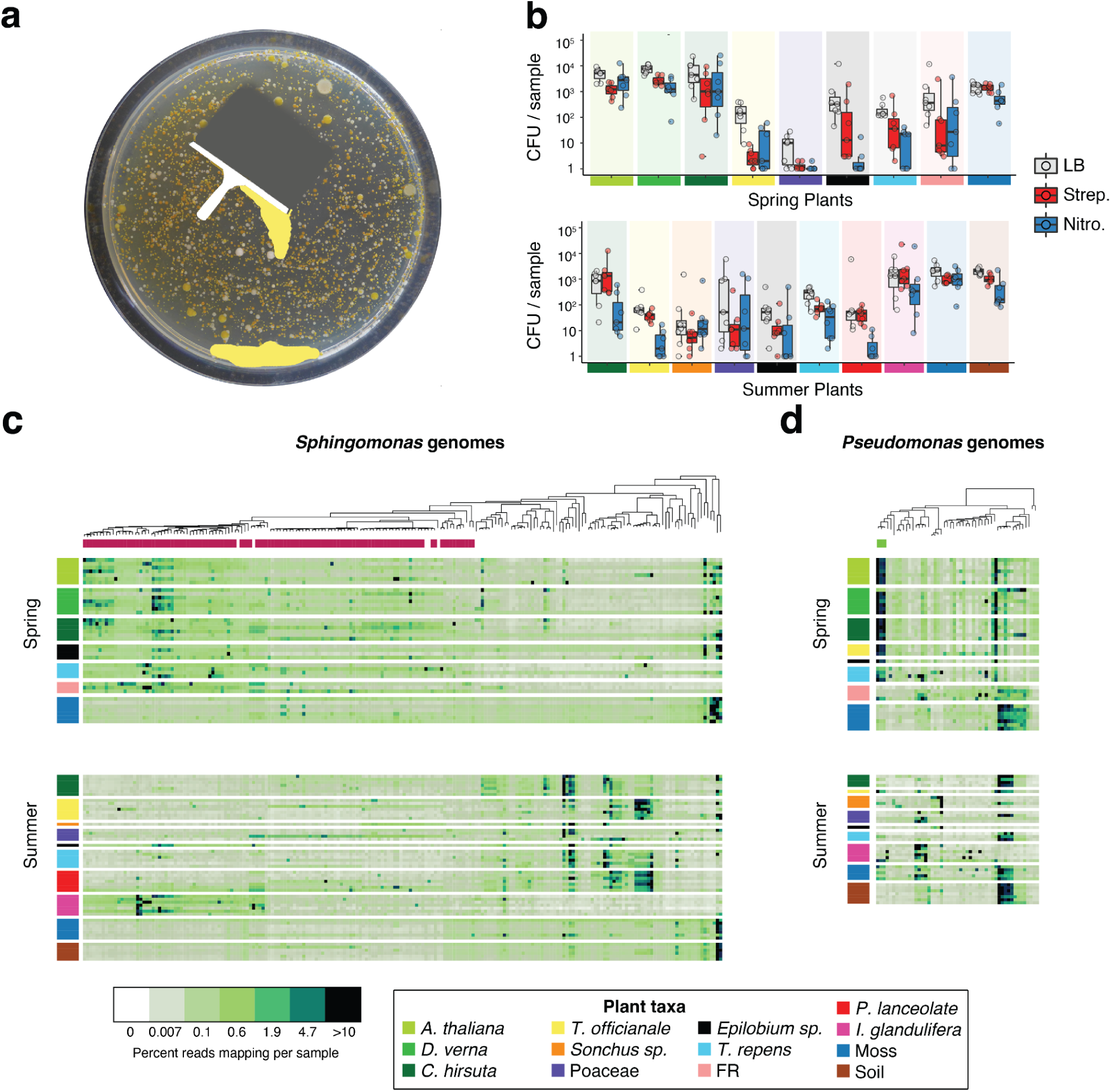
**Metagenomics of cultured bacteria reveals strain sharing across plant species. a**, Cartoon demonstrating the process of generating and harvesting bulk cultures for plant-free metagenomics. **b**, Colony forming units for each plant host species (see color legend, bottom left) on medium containing no antibiotic (LB), *Sphingomonas*-selective streptomycin (Strep) medium, or *Pseudomonas*-selective nitrofurantoin (Nitro) medium. Each sample represents about 4.5 mg fresh weight of original leaf material. **c**, Heatmap showing fourth-root transformed reads from each *Sphingomonas* bulk culture metagenome (rows, colors from plant host taxa shown in the legend below) that map to a *Sphingomonas* local reference genome (columns), for the spring collection (top) and the summer collection (bottom). The genetic relatedness of the local bacterial reference genomes is shown by a maximum likelihood (ML) tree above the heatmap. Those reference genomes belonging to SphASV1 are indicated under the ML tree in magenta. Darker colors in the heatmap correspond to genomes attracting a greater fraction of reads in the mapping process. **d**, Same as **(c)**, but showing *Pseudomonas* bulk culture mapping to *Pseudomonas* reference genomes. Reference genomes belonging to PseASV1 are indicated under the ML tree in green.

To harvest the bacteria, we flooded the plates with PBS and then scraped all colonies from a plate with a flame-sterilized razor blade off into a single bacterial pool, from which we prepared whole metagenome shotgun libraries (Fig. 5a). This culture-dependent metagenomic strategy (*71*) allowed us to investigate endophyte-enriched bacteria while completely avoiding plant DNA. After sequencing the metagenomes using paired end 2×150 bp Illumina reads, we mapped the reads to a comprehensive reference database including all of our local genomes plus selected publicly-available genomes (*54, 72*) (Methods, Extended Data Fig. 8). Because some colonies did not belong to our targeted genera, we also included in our reference database “decoy” genomes of common plant-associated bacteria to capture reads from these “contaminant” strains (Extended Data Fig. 9). A total of 165 metagenomes (91 *Sphingomonas* + 74 *Pseudomonas*) passed our quality thresholds. For both genera, there was a clear shift in dominant strains across seasons, including for those host plants that were alive and were sampled at both timepoints (Fig. 5c, 5d). This seasonal shift was also apparent in the *Sphingomonas* and *Pseudomonas* amplicon data (Fig. 4a, 4b). Finally, many metagenome reads from diverse plant samples mapped to the same strains, in agreement with them being widely shared across plants. In particular, this could be observed for strains associated with the most abundant 16S rDNA sequences, SphASV1 and PseASV1. The relative representation of both SphASV1 and PseASV1 isolates in the summer soil samples was low compared to that in plants, suggesting that as the new cohort of *A. thaliana* germinates in the following fall, it may be more likely that *A. thaliana* seedlings acquire these strains from nearby plants rather than from the soil.

### *Pseudomonas* are much stronger leaf saprophytes than *Sphingomonas*

We consistently noticed a slower growth rate for *Sphingomonas* than for *Pseudomonas*, both on selective media and when we streaked individual strains on non-selective medium, with colonies of SphASV1 often appearing only up to three days later than PseASV1 colonies. This surprised us, because SphASV1 seemed to be as successful as PseASV1 in establishing substantial population sizes in leaves of wild plants. We hypothesized that LB medium might advantage *Pseudomonas*, and growth of the bacteria in the more complex nutrient and chemical milieu supplied by plants might result in more equal performance between the genera. To test this, we collected wild leaves of both *A. thaliana* and *Brassica napus* from a site in Kusterdingen, Germany in May 2021. For both *A. thaliana* and *B. napus* leaves, PseASV1 and SphASV1 were present and highly abundant in all starting leaf material, consistent with other collections. We macerated a subset of the leaves with a mortar and pestle, and then compared the growth of bacteria over the next two days within macerated leaves and within detached but un-macerated control leaves (Fig. 6). We incubated all plant material in sealed Petri plates in 16 hours of light, at a constant 16°C, which is close to average daytime spring temperatures at the site of our collections. At each timepoint, we quantified bacterial relative abundances with 16S rDNA amplicon sequencing to detect all genera, and we cultivated and counted *Pseudomonas* CFUs, which are easier to exclusively isolate and quicker to grow than *Sphingomonas*, to estimate changes in absolute bacterial abundances.

**Fig. 6.**
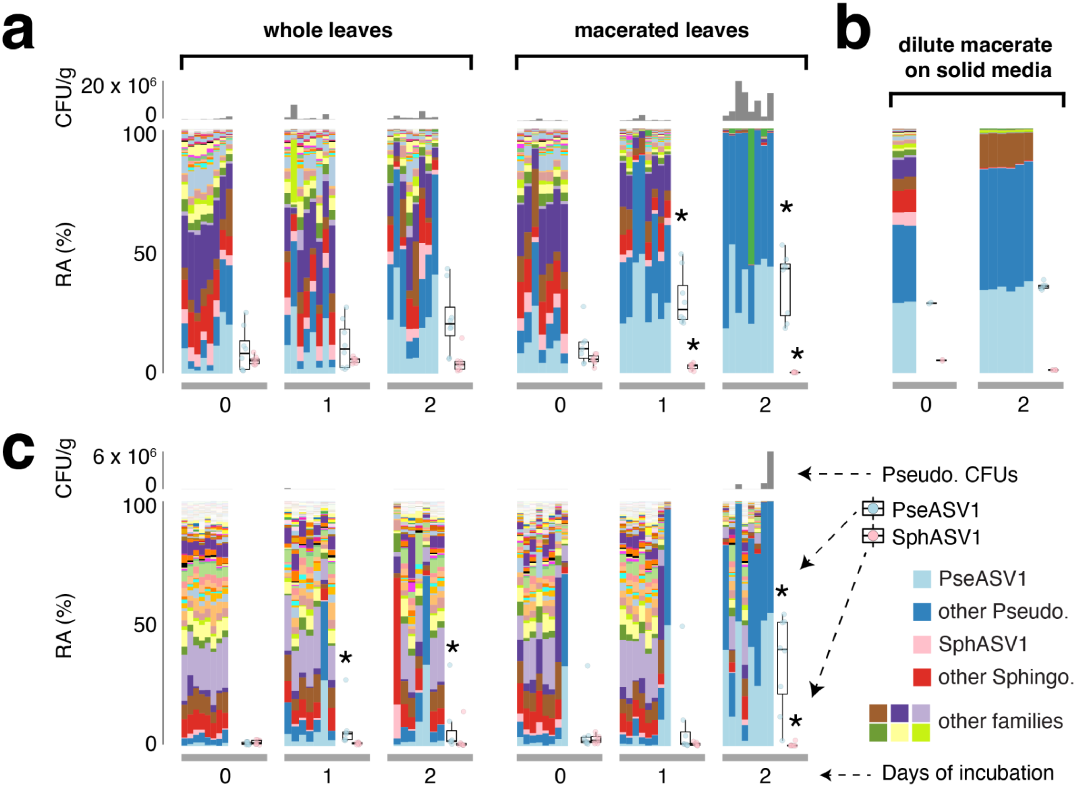
**Contrasting *Pseudomonas* and *Sphingomonas* growth in whole vs macerated leaves. a**, Bacteria growing in wild *A. thaliana* whole leaves (left) or macerated leaves (right) at 0, 1, and 2 days post harvesting. The layout for all panels is as diagrammed on the right of panel **(c)**, with *Pseudomonas* CFU/g shown as a grey column chart (top), the relative abundance (RA) of different bacterial families shown as a stacked barplot colored according to the legend, and with the relative abundances of PseASV1 and SphASV1 for each timepoint summarized as box plots. An asterisk (*) represents a significant difference from 0 days of incubation (*p*<0.05 according to a Mann-Whitney U-test). **b**, Similar to **(a)**, but with bacteria from macerated *A. thaliana* leaves at the time of harvest and after 2 days of incubation on LB plates. Pseudomonas CFUs were not counted **c**, Same as **(a)**, but with bacteria in wild *B. napus* leaves instead of *A. thaliana*.

PseASV1 proliferated to represent a strong majority of bacteria in most *A. thaliana* and *B. napus* leaf macerates (Fig. 6a and 6c). While PseASV1 also increased its relative abundance in detached whole leaves of both plant species over the same timeframe, the magnitude was much less pronounced than in macerates. We also spotted liquid collected from *A. thaliana* leaf macerate onto solid LB agar and incubated it in the same conditions, allowed all cultivable bacteria to compete with each other on the plate, and examined the bacterial population after 2 days (Fig. 6b). Although some changes in relative abundance of 16S rDNA sequences from bacteria on LB media relative to the starting leaves were likely due to the loss of uncultivable bacteria that cannot grow on LB, our interest was in cultivable PseASV1- and SphASV1-associated strains. As in macerated plant tissue, PseASV1 strains on LB markedly increased their abundances in the community after 2 days, while the relative abundance of SphASV1 strains and indeed all Sphingomonadaceae decreased to essentially zero (Fig. 6b). These results strongly suggest that PseASV1 thrives in dead or dying leaves and furthermore on rich non-living substrates, with the intriguing corollary being that *Sphingomonas* depends on healthy leaves to maintain its competitive edge.

## Discussion

Most microbes that live in or on multicellular organisms do not have an obligate relationship with any specific multicellular species, but rather are better adapted to some, which are called hosts, than to others, which are called non-hosts (*73, 74*). We sought to use genomic and metagenomic techniques to characterize, at the strain level, the extent of host specialization for the two bacterial genera that are locally the most abundant in the phyllosphere of the host plant *A. thaliana*. We found a higher genomic diversity of *Sphingomonas*, which generally reaches very similar population sizes in the leaves of our plant populations, than for *Pseudomonas*, which has very different population sizes in different individuals. The high genomic diversity of *Sphingomonas* held true even among those sharing the same 16S rDNA sequence, highlighting the importance of strain-resolved techniques also in wild ecosystems. To this end we employed direct culturing and sequencing of individual isolates, as well as bulk-culture metagenomics (Fig. 5). While culturing is less quantitative and potentially less inclusive than direct sequencing, genetically diverse *Sphingomonas* and *Pseudomonas* can be cultured with low bias (Fig. 1f, 1g), making this a powerful technique to further reveal colonization patterns for strains in these genera.

Our previous work characterizing PseASV1/*Pv-ATUE5* suggested that much of its standing genetic variation predates *A. thaliana* colonizing SW Germany (*19*), and we had puzzled how ancient variants are apparently able to continue to coinfect the entire population in this pathosystem because in agricultural systems epidemics are typically monomorphic with a rapid turnover of pathogenic isolates within a few years (*75, 76*). The present study begins to answer this question. First, PseASV1/*Pv-ATUE5* strains efficiently colonize other local plant species besides *A. thaliana*, especially other Brassicaceae. The same *Pseudomonas* strains observed on *A. thaliana* also persist through the summer, living on other hosts when there are no alive *A. thaliana* plants, implying that their performance on these additional hosts may be equally, if not more important to their long-term success than their seasonal exploitation of *A. thaliana* as a host. Currently, we do not know how important different hosts are for different subclades of PseASV1/*Pv-ATUE5*, but with a variety of host species as well as intraspecific genetic heterogeneity in each host, it may be difficult for any one strain to prevail and dominate the entire *Pseudomonas* population – matching a scenario of diffuse interactions, as proposed before as an explanation for non-matching distributions of a specific *Pseudomonas syringae* effector and the *A. thaliana* resistance locus that detects it (*77*). In contrast, in a crop monoculture, genetically-identical hosts provide more consistent and uniform challenges for a pathogen, making it more likely for a single strain to become dominant. A second explanation for the lack of a single emergent strain is that the pathogenic nature of PseASV1/*Pv-ATUE5* is strongly context-dependent, and apparently more so than for other *Pseudomonas* pathogens such as *Pst* DC3000. In an agar plate system, disease caused by PseASV1/*Pv-ATUE5* was much more severe than that of *Pst* DC3000, but in potting soil, PseASV1/*Pv-ATUE5* failed to cause significant disease symptoms and grew much more slowly than *Pst* DC3000. Consistent with fitness effects of PseASV1/*Pv-ATUE5* being greatly modulated by the environment, we saw few disease symptoms on field-collected plants at the time of collection, regardless of their *Pseudomonas* load.

Compared to PseASV1/*Pv-ATUE5*, *Sphingomonas* strains that shared the most abundant V3V4 *Sphingomonas* ASV, SphASV1, were more than three times as diverse by average nucleotide identity in their core genomes. SphASV1 isolates differed in type 4 and type 6 secretion system presence, anoxygenic photosynthetic ability, plasmid presence, and in other unidentified features that are likely to affect host colonization potential or intermicrobial competition. The plasmid count, up to three per strain in our closed genomes, suggests that the *Sphingomonas* genetic tool kit may be highly modular. We observed SphASV1 in diverse 16S rDNA datasets worldwide, including high abundances on *Boechera stricta* in North America (*14*), and while the extent of genome variation among SphASV1 strains in SW Germany makes it difficult to predict genomic features from 16S rDNA, it will be highly interesting to determine the global diversity in this group of *Sphingomonas* strains.

This study also demonstrated that apart from SphASV1, the genus *Sphingomonas* is not only abundant, but consistently so across plants (Fig. 1). It colonizes diverse plants at our field site to high levels (Extended Data Fig. 3), a result consistent with reports of high abundances on leaves of diverse plants such as maize (*15*), poplar (*11, 12*), and other species (*11*). The different colonization patterns of the genera *Pseudomonas* and *Sphingomonas* provide clues about their lifestyle strategies. The fact that *Pseudomonas* populations vary greatly in size between plants, occasionally reaching high abundances as homogenous blooms of PseASV1/*Pv-ATUE5* (*19*), suggests that different strains may compete for resources and share the same niche, making stable coexistence within a plant less likely. In contrast, *Sphingomonas* loads varied little across plants, regardless of the load of other bacteria (Fig. 1a), and the diversity of *Sphingomonas* ASVs in each plant was also higher. The more balanced coexistence of different *Sphingomonas* strains may mean that each occupies a different microniche. To what extent these bacteria inhabit spatially-distinct parts of the leaf or grow as biofilms is unknown; future direct visualization techniques will help resolve this.

Some *Sphingomonas* can project against pathogenic *Pseudomonas* and other bacterial and fungal pathogens, and they also affect overall microbial community structure (*6, 33*). This may be in part because their metabolic needs substantially reduce the availability of substrates for other phyllosphere microbes (*35*). However, in at least one known case, a *Sphingomonas* strain secretes an extracellular small molecule that attenuates the virulence of a bacterial pathogen (*22*). Further, the plant protective *S. melonis* Fr1 induces transcriptional changes in *A. thaliana* and protection depends on the presence of a plant immune receptor (*62*). These examples illustrate that protective mechanisms go beyond substrate competition. Although we estimate that in our culture collection only a minority of *Sphingomonas* strains have protective ability against PseASV1/*Pv-ATUE5* or *Pst* DC3000, the high bacterial titres we inoculated with and the gnotobiotic environment in which we observed the strongest effects may limit transferability of our results to the field, especially as the most protective local *Sphingomonas* isolate had a 16S rDNA sequence that was not among the more abundant on field plants. The high diversity of *Sphingomonas* genomes, the presence of secretion systems often involved in biotic interactions, and the fact that we observed some antagonism suggests there is more to be discovered. The overall implications of *Sphingomonas* populations on leaves will be important to understand, both for their direct impacts on plant recognition and tolerance of non-pathogens, and also for their role in structuring wild microbial communities.

The evolutionary pressures on these generalist bacteria are many (*78*), involving at a minimum multiple host plants with different immune system activities as well as free-living phases. Our final experiment demonstrating PseASV1-associated strains’ growth advantage over SphASV1-associated strains in detached leaves, macerated leaves, and solid media hints at the fact that very different growth strategies drive dominance in the phyllosphere. *Sphingomonas* grows more slowly and appears to thrive in healthy leaves, perhaps because of a higher investment in defense against the plant, and we speculate that investing in plant-beneficial features may be part of a strategy to co-opt the plant’s immune system to help prevent opportunists or saprophytes from outcompeting it. In contrast, *Pseudomonas* strains quickly overtake weakened and dead leaves, and while they may survive on healthy leaves, occasionally weakening the plant immune system can promote greater populations and bacterial spread.

## Materials and Methods

### Community plant collection and lysate glycerol stock preparation

For the community analyses, plants were harvested at Eyach (community of Starzach, 48°26’46.0"N 8°47’02.4"E) in Germany (Extended Data Fig. 1). The spring visit occurred on the April 20, 2018, and the summer visit occurred on the September 14, 2018. For each plant species, we pooled together entire leaves from at least 6 independent plants per sample, and collected 7 such independent samples. After bringing samples back to the lab, we surface-sanitized them in 70% ethanol for 45-60 s (*41*), ground the fresh tissue using a sterile mortar and pestle in a volume of PBS proportional to the sample’s fresh weight (7.4 mL PBS per gram of tissue), and mixed the resulting lysate with glycerol to make -80°C freezer stocks with a final glycerol concentration of ∼27%, suitable both for direct DNA extraction and culturing.

### Collection of *Sphingomonas* isolates

Most *Sphingomonas* isolates (Extended Data Table 2) were cultured from frozen *A. thaliana* lysates in glycerol that had originally been prepared stored on December 11, 2015 and March 3, 2016 (*19*). Briefly, we processed two leaves each per *A. thaliana* rosette. Each leaf was washed in 75% EtOH for 3-5 s, the ethanol was allowed to evaporate in a sterile hood, and the leaf was ground in 10 mM MgSO_4_ before being mixed with glycerol to a final concentration of 15%-30%. The glycerol stocks were stored at -80°C. From each sample, 75 mL of lysate was plated on large petri plates (145 x 20 mm, Greiner Bio One International GmbH, Frickenhausen, Germany) containing low sodium Luria broth (LB, 10 gL peptone, 5 g/L yeast extract, 5 g/L NaCl) and 1.5% agar supplemented with 100 µg/mL cycloheximide to suppress fungi (SERVA Electrophoresis GmbH) and 100 µg/mL streptomycin (Thermo Fisher Scientific, Karlsruhe, Germany). Plates were incubated at room temperature for 5 to 10 days, then stored at 4°C until processing. The remaining *Sphingomonas* isolates from pooled *A. thaliana* and other plant species were recovered similarly by plating lysates on selective media (see section “Bulk culture and additional cultured isolates”).

Up to 16 colonies per plate were randomly picked and transferred to 24-well plates (CELLSTAR, Greiner Bio One International GmbH, Frickenhausen, Germany) containing the same selective LB agar, and incubated for 5-10 additional days as necessary to create a small bacterial lawn. All bacteria were cultivated on agar, as we observed that many grew poorly in liquid media. Bacteria were then scraped from the agar with a sterile loop and transferred to a 2 mL screw cap tube (Starstedt, Nümbrecht, Germany) filled with 300 ml of PBS (we later switched to 300 mL of liquid LB media instead of PBS, which improved survival). Three sterile glass balls (Æ 2.85-3.45 mm or Æ 0.25-0.5 mm) (Carl Roth GmbH, Karlsruhe, Germany) were added to each screw cap to help dissociate (but not lyse) clumped bacterial cells, and tubes were shaken in a FastPrep 24™ 5G homogeniser (MP Biomedicals, Eschwege, Germany) at speed 4.0 m/s for 10 seconds. From the mixed cell suspension, 250 mL was stored at -80°C in ∼27% glycerol.

### Sphingomonas DNA isolation for short read sequencing

The remaining 50 mL was mixed with 150 mL of sterile PBS and transferred to a semi-skirted 96-well PCR microplates (Axygen™) for DNA extraction. Bacterial cell suspensions were lysed by incubation with lysozyme (100 µg/mL final concentration) at 37°C for 30 min followed by incubation with sodium dodecyl sulfate SDS (1.5% final concentration) at 56°C for 1-2 h. Next, ⅓ volume (66 μL) of 5 M potassium acetate (CH_3_COOK) was added to each well to precipitate the SDS and other cytosolic components. The 96-well plates were centrifuged at maximum speed for 20 min to pellet the precipitate, and the supernatant transferred to a new 96-well plate for genomic DNA purification using Solid Phase Reversible iImmobilization (SPRI) magnetic beads (*79*). Briefly, home-made SPRI bead mix adapted from (*79*) was thoroughly mixed with samples at a ratio of 0.6:1 bead to sample ratio and incubated for 15-20 min. The plates were placed on a magnet for 5 min and the supernatant was removed. Following two 80% EtOH washes, the beads were air-dried and resuspended in 50 μL elution buffer (EB, 10 mM Tris, pH 8.0). After an overnight incubation at 4°C, the plates were placed in the magnet and the elution buffer containing the DNA was transferred into a new 96-well PCR plate.

### Bulk culture and additional cultured isolates

For *Pseudomonas* and *Sphingomonas* bulk culture analysis, approximately 50 mg of plant lysates collected from Eyach in 2018 was scooped from the frozen glycerol stocks at -80°C, and after thawing, 50 μL of the thawed lysate was pipetted onto 2% agar LB plates (200 mm diameter), using an additional 150 μL of sterile PBS to aid in spreading. The LB medium was supplemented with 100 μg/mL cycloheximide and 100 μg/mL streptomycin for *Sphingomonas* bulk culturing, and 100 μg/mL cycloheximide and 100 μg/mL nitrofurantoin (Sigma-Aldrich, Steinheim am Albuch, Germany) for *Pseudomonas* bulk culturing (*19*). Plates were incubated at room temperature for 5 days for *Sphingomonas*, and for 1.5 to 2 days for *Pseudomonas*.

Individual colonies from selected plates were randomly picked to 24-well plates, as described above. To harvest the remaining bacterial colonies in bulk, plates were soaked for 5 min with 4 mL of PBS, and the surface was scraped with a flame-sterilized razor blade to loosen adherent bacteria. The plate was tilted to form a pool of PBS at the lower end, and the scraped bacteria were mixed into the PBS pool by pipetting up and down. An appropriate aliquot of the mixed suspension was then transferred to a 2 mL centrifuge tube, such that the eventual pellet would extend a maximum of 5 mm from the base of the tube (which approaches an upper limit for efficient and uniform DNA extraction using our methods).

### DNA extraction from bulk culture bacterial pellets and plant lysates

For bulk culture bacterial pellets, the final pellets were resuspended in at least 750 μL of DNA lysis buffer containing 10 mM Tris pH 8.0, 10 mM EDTA, 100 mM NaCl, and 1.5% SDS. Especially large pellets (greater than 5 mm from the bottom of the tube) were suspended in proportionally greater volumes of buffer to ensure an efficient lysis, and 750 μL was used for lysis. For DNA extraction from the plant lysates in glycerol, 400 uL of glycerol lysates was mixed with 400 μL of the DNA lysis buffer described in the previous sentence, but with 3% SDS instead of 1.5% SDS to yield a final SDS concentration of 1.5%. The suspensions were pipetted to a screw cap tube containing ∼0.5 mL sterile garnet rocks (Bio Spec Products Inc., Bartlesville, USA) and homogenized in a FastPrep 24™ at speed 6.0 m/s for 1 min. The tubes were next centrifuged at 10000 x g for 5 min, and the supernatant (about 600 μL) mixed with 200 μL sterile 5 M potassium acetate in a 1 mL 96-well plate (Ritter Riplate® 43001-0016, Schwabmünchen, Germany) to precipitate the SDS. The plates were spun at 5000 x g for 10 min and the supernatant transferred to a new 1 mL deepwell plate. The resulting supernatant was centrifuged a second time to clear out remaining plant material and precipitate. Finally, 360 μL SPRI beads were added to 600 μL of the supernatant. After mixing and incubating on a 96-Well Magnet Type A (Qiagen, Hilden, Germany) the beads were cleaned with 80% ethanol and DNA was eluted in 100 μL of EB.

### 16S rDNA V3-V4 amplicon sequencing

The 16S rDNA V3-V4 region was amplified using the 2-step protocol described for V4 amplicons in (*10*), with the exception that the forward PCR primer was 341F (Extended Data Table 1), and due to this different forward primer, the annealing step in the first PCR was done at 55 °C instead of 50 °C in (*10*). In addition, because the amplicons were longer than the V4 amplicons in (*10*), the libraries were sequenced on MiSeq instrument (Illumina) with a V3 2 × 300 bp reagent kit instead of a V2 2 × 250 bp reagent kit. This allowed overlap and assembly of the forward and reverse reads. The frameshifts built into the primers used in the first PCR made the addition of Illumina PhiX control library to increase sequence diversity unnecessary [27], and it also enabled barcoding of more samples. This is because half of the samples from the first PCR were amplified with 341F frameshifts 1, 3, and 5 paired with 806R frameshifts 2, 4, and 6. The other half of the samples from the first PCR paired 341F frameshits 2, 4, and 6 with 806R reverse frameshifts 1, 3, and 5. The first PCR used 29 cycles and the second PCR used 6 cycles for a total of 35 cycles. We denoised the sequences into amplicon sequence variants (ASVs) using USEARCH (*80*), and classified the ASVs to bacterial taxa against the RDP database (*81*).

### Bacterial whole-genome sequencing and bulk culture metagenome sequencing with short reads

DNA from each bacterial isolate, or from each metagenome, was quantified by Quant-iT™ PicoGreen™ dsDNA (Invitrogen™) in a Magellan Infinite 200 PRO plate reader (Tecan Trading AG, Switzerland) and diluted and normalised to 0.5 ng/μl as a prior step to library construction. Bacterial DNA libraries were constructed using an adapted Nextera™ protocol for small volumes (*19, 82*). In brief, 2.5 ng of DNA was sheared using Nextera Tn5 transposase (Illumina, Inc), and Nextera sequencing adapters were added though 12 cycles of PCR as previously described (*19*). An aliquot of each library was run in an agarose gel for quality control, and the remainder of the library was purified using SPRI beads to remove primers at a ratio of 1.5:1 beads to PCR product. The clean DNA was eluted in 40 μL of EB and the concentration of the final product was quantified with PicoGreen. Libraries were pooled in an equimolar ratio. To increase the concentration of the pool to enable further size selection procedures, the pooled library was concentrated by first precipitating the DNA by mixing with it with an equal volume of a solution containing 1 part sodium acetate and 8 parts isopropanol, passing the solution over an EconoSpin Mini Spin Columns (Epoch Life Science Inc., Missouri City, USA), washing the column twice in 70% ethanol, and eluting in 50 μL EB. The resulting multiplexed, concentrated library molecules were size-selected to keep fragments between 350 and 700 bp using a 1.5% cassette in a BluePippin instrument (Sage Science Inc., Beverly, USA). After adjusting the concentration of each size-selected pool to 2.5 nM, the DNA was sequenced with 2 ×150 bp paired-end reads on a HiSeq 3000 instrument (Illumina inc, San Diego, USA). Metadata regarding sequenced genome and metagenome samples can be found in Extended Data Table 2 .

### Short-read genome assembly and annotation

Genomes were assembled using SPAdes genome assembler version 3.11.0 correcting for mismatches and short indels (--careful) and k-mer sizes of 21, 33, 55, 77, 99 and 127 (-k) (*83*). Draft genomes were corrected using Pilon version 1.20 with standard parameters (*84*). Annotation of bacterial genomes was accomplished using Prokka version 1.12 using the --compliant parameter (*47*). Coverage and N50 statistics were measured using custom scripts, including the N50.sh script from the GitHub repository of Henk den Bakker (*85*). The completeness of the genome was assessed using BUSCO version 2.0 (*46*) selecting proteobacteria lineage and the gene set (proteins) assessment (-m prot).

### Production and assembly of closed Sphingomonas genomes

Assembly of closed *Sphingomonas* genomes required a separate DNA preparation and library production for sequencing on an Oxford Nanopore MinION instrument. First, 12 *Sphingomonas* isolates were each bulked on 15 × 15 cm^2^ LB plates with 100 µg / mL streptomycin. After 2-5 days, depending on the growth rate of each strain, the bacterial lawn was harvested, and ∼100 mg of bacteria was mixed with 800 µL lysis buffer, and DNA was extracted and purified using the homemade SPRI beads exactly as described above for bulk culture lysates, including the harsh bead beating. We noticed that bead beating did not shear DNA below 10 kb, and fragments greater than approximately 7 kb are sufficiently long to span repeated regions in bacterial genomes (*86*). The harsh bead beating during the lysis step naturally sheared the DNA to approximately 15-20 kb, and the 0.6 : 1.0 SPRI bead cleanup removed most of the smaller fragments, so no additional shearing or size selection was performed. DNA was eluted from the SPRI beads in 400 μL EB. Because some DNA extracts remained discolored or viscous following the SPRI purification, the DNA was further purified by mixing with it with an equal volume of chloroform. The aqueous phase was collected and the HMW DNA was precipitated by mixing it with an equal volume of a solution containing 1 part sodium acetate and 8 parts isopropanol. Precipitated DNA was pelleted by centrifugation at 20,000 x g for 5 minutes. The pellet was washed twice in 70% ethanol and eluted again in 400 μL EB.

Pure DNA was prepared for sequencing on the Oxford Nanopore MinION (Oxford Nanopore Technologies, Oxford, UK) using the manufacturer-recommended protocol “1D Native barcoding genomic DNA” using the SQK-LSK109 kit with barcode expansions NBC104 and NBC114 for 24 samples. Briefly, DNA concentration was adjusted to 1 μg of DNA diluted in 49 μL of NFW water (Ambion). From this 48 μL were treated with NEBNext reagents for FFPE and end repair as well as dA-tailing. The end-repaired libraries were cleaned with Agencourt AMPure XP beads at a 1:1 ratio and eluted in 25 μL in a 1.5 mL DNA LoBind tube (Eppendorf AG, Hamburg, Germany). Native barcodes were attached by ligation, the solution was again cleaned with AMPure beads at a 1:1 ratio, and concentration of all barcoded-added samples were measured using a Qubit (Thermo Fisher Scientific Inc., Waltham, USA) fluorometer and pooled at equimolar ratios for a total of 700 ng. Finally, sequencing adapters were added to the pool by ligation. LFB was selected for washing in the final clean-up with AMPure beads. The library was prepped and loaded into an FLO-MIN106 RevD R9.4.1 flow cell following the manufacturer’s instructions and sequencing was run for 24 h.

To assemble closed *Sphingomonas* genomes, Guppy package version 3.0.3 (https://nanoporetech.com/) was first employed to perform initial base calling to produce raw read and quality assessment in the FASTQ format as recommended by (*87*). Next, the samples were demultiplexed using qcat. Draft contigs were then assembled *de novo* by using mini_assemble which is part of the Pomoxis toolkit (version 0.3.6, https://github.com/nanoporetech/pomoxis) using default parameters for de-novo assembly employing miniasm (*88*) and four rounds of long-reads-based polishing with minimap2 and Racon. Each of the genomic consensus assemblies was further improved by additional 4 rounds of long-reads-based polishing with Racon followed by one additional round using medaka_consensus (package version 0.6.5, https://github.com/nanoporetech/medaka). To enhance base corrections, an additional polishing step using high-quality Illumina short reads from the same *Sphingomonas* strains was used. First, the short reads were mapped onto the assembled *Sphingomonas* genome, using Burrows-Wheeler Alignment (BWA) with BWA-MEM (*89*). Next, these mapped reads were used to correct the assembly using Pilon version 1.23 with default parameters (*84*). Assessment of genome completeness and annotations were performed in the same way as short-read draft genomes. Metadata regarding sequenced closed genomes can be found in Extended Data Table 2 .

### Pan-genome and phylogenetics analysis

The panX pan-genome pipeline (*53*) was used to assign orthology clusters and construct the phylogenetic tree, taking as input the Genbank format files (.gbk) from Prokka (previous section). The following parameters were used for the panX analysis: the divide-and-conquer algorithm (-dmdc), a size of 50 strains per subset to run DIAMOND (-dcs 50) (*90*), and a soft core genome cutoff of 70% that includes all genes present in >70% of the strains as part of the core genome (-cg 0.7). Genomes included in each panX run are indicated in Extended Data Table 2.

### Processing bulk culture metagenomic reads

A first quality control of raw sequencing data was performed using Skewer version 0.2.2 (*91*) to trim raw reads and remove highly degenerative reads (-n) or reads shorter than 20 bp (-l 20). For those samples with a total yield of at least 500 Mb, the filtered reads were mapped with BWA-MEM (*89*) using standard parameters against a custom-made reference database (described in the next section). Those reads mapping with a quality score of 30 or higher were output in a BAM file using SAMtools (*92*). Duplicated reads were removed from BAM files using the MarkDuplicates command in Picard tools (http://broadinstitute.github.io/picard/) version 2.0.1 using default parameters. SAMtools -stats and -fastq commands were used to retrieve BWA-MEM mapping statistics and convert the BAM files back to FASTQ files, respectively (Extended Data Fig. 8).

### Custom local reference database

Metagenome reads from the *Sphingomonas* or *Pseudomonas* bulk cultures were mapped to a three part custom-made reference database of bacterial genomes. First, “Decoy” genomes contained bacteria from other genera, and were used to help classify reads from contaminant bacteria. The collection of Decoy genomes was made by downloading the *A. thaliana* phyllosphere, root, and soil isolates from (*72*) and removing *Pseudomonas* when mapping *Sphingomonas,* or removing *Sphingomonas* when mapping *Pseudomonas.* Three different genome sets were combined with distinctive headings to allow later recovery of mapped reads:

<u>DECOY</u>: A set of bacterial genomes (∼430 genomes) found in *A. thaliana* phyllosphere, roots, and soil surrounding the plants. Published in (*72*). Decoy genomes have the mission to capture contaminant reads.

<u>REFSEQ</u>: A set of bacterial genomes of *Pseudomonas* or *Sphingomonas* found in the NCBI Reference Sequence Database (*54*).

<u>LOCAL</u>: A set of bacterial genomes of interest sequenced in-house. This includes 165 genomes representative of local *Pseudomonas* isolates sequenced by (*19*) and all the single *Sphingomonas* and *Pseudomonas* genomes from this study.

### Genome similarity comparisons

Briefly, to generate similarity matrices for *Sphingomonas* and/or *Pseudomonas* genome comparisons using MASH (version 2.1)(*49*), we used the formula 100 ⨉ (1-MD), where MD is the MASH distance, to convert MASH distances to similarity scores. To calculate average nucleotide identity (ANI), we used FastANI (*50*) For similarity matrices Heatmaps of similarity scores were illustrated using the function “heatmap.2” in the R package “gplots” (*93*).

### Inoculation, infection, and phenotyping in 24-well plates

<u>Plant cultivation:</u> Each well of a 24-well plate (Greiner) was filled with 1.5 mL of 1% agar (Duchefa) containing ½ strength Murashige-Skoog (MS) medium with MES buffer. *A. thaliana* seeds (accession Ey15-2, CS76309) were surface-sterilized by submerging in 70% EtOH with 0.01% Triton X-100 for 1 min, then submerging in 10% household bleach solution for 12 min, and finally washing three times with sterile water. The seeds were then stratified at 4°C for 3 days in water, and then were pipetted onto the agar (1 seed/well). Excess water after distributing seeds was removed by pipetting.

<u>Inoculation and infection:</u> Freshly-planted, ungerminated seeds were inoculated with 4 µL of 10 mM MgCl₂ or with *Sphingomonas* sp. suspended in the same amount of buffer to an optical density at 600 nm (OD_600nm_) of 0.5. The *Sphingomonas* colonies had been previously cultivated for 5 days on selective LB agar plates with 100 µg/mL streptomycin, were scraped from the plates with a sterile loop, and were washed twice by centrifugation and resuspension in MgCl₂ to remove residual antibiotics. Inoculated seeds were germinated and seedlings were cultivated for 10 days in growth chambers at 21°C with 16 hours of light. The seedlings were then challenged with 100 µL of 10 mM MgCl₂ or with *Pseudomonas* suspended in the same amount of buffer to OD_600nm_ = 0.01. The bacteria were drip-inoculated by pipette to the center of each rosette. Plates were sealed with Parafilm and returned to the growth chamber for 7 days.

<u>Plant phenotyping:</u> On 0, 2, 4, and 7 days post inoculation, rosettes were imaged in the plates with a custom procedure to eliminate glare. Briefly, an opaque box was filled with a LED light source and covered with a sturdy translucent paper surface to diffuse the light. Each plate was placed on top of the paper in a defined position, and the backlighting allowed imaging from above without removing the lids. Pictures were taken using a LUMIX DMC-TZ71 digital camera (Panasonic Co., Osaka, Japan) without flash. Images were processed similarly to that described in (*94*). Briefly a predefined mask was used to extract each plant in the image, and automatic segmentation based on pixel color was applied to recognize plant leaves from background. The leaf area of each plant was then calculated based on the segmented plant images.

### Inoculation, infection, phenotyping, and hamPCR of plants on potting soil

<u>Plant cultivation:</u> *A. thaliana* seeds were surface-sterilized, stratified for 4 days in sterile water at 4℃, and sowed on potting soil (CL T Topferde; www.einheitserde.de) in 7 cm pots (PL 2832/20, Pöppelmann GmbH, Lohne, Germany). Seedlings were germinated and cultivated under short day (8 hr light) growing conditions at 23°C and 65% relative humidity, illuminated by cool white fluorescent light of 125 to 175 μmol×m^-2^×s^-1^.

<u>Inoculation and infection:</u> Two weeks after sowing, seedling leaves were sprayed ad- and ab- axially with *Sphingomonas* strains cultivated identically as for inoculation in 24-well plates as described above, but resuspended in 10 mM MgCl_2_ buffer to a concentration of OD_600nm_ = 1.0. Plants were also sprayed with heat-killed (boiled) *Sphingomonas* prepared by mixing equal parts of all strains after they had been resuspended at OD_600nm_ = 1.0 in MgCl_2_ and boiling the resulting solution for 10 minutes. Humidity domes were kept on the flats for 48 hours. On the fourth day following *Sphingomonas* treatment (96 h), plants were sprayed with *Pseudomonas* strains also cultivated identically as for 24-well plates, but resuspended in 10 mM MgCl_2_ buffer to a concentration of OD_600nm_ = 1.0. A heat-killed *Pseudomonas* mix was also prepared, as described above. Prior to spray-inoculating *Pseudomonas,* the surfactant Silwet L-77 added at 0.04% v/v following the protocol in (*63*). Humidity domes were kept on the flats for 72 hours.

<u>Plant phenotyping:</u> Five days (120 h) post inoculation, overhead images of pots arranged in the flat were taken from a height of ∼ 1.5 M with a LUMIX DMC-TZ71 digital camera. Because the position of each pot in the flat and the position of each plant in the pot was not fixed in the overhead images, identically-sized squares containing each plant were manually cropped from each image and arranged into a montage for each flat using object alignment functions in Adobe Illustrator. The montages were then processed programmatically as described in the previous section by using a predefined mask to extract each plant in the image and counting green pixels (Extended Data Table 3).

<u>Confirmation of viable bacteria and DNA extraction:</u> Following plant phenotyping, whole seedlings were harvested to 2 mL screw-cap tubes (Type I, Sarstedt AG, Nümbrecht, Germany) using flame-sterilized tweezers and scissors, and kept on ice. First, one 5 mm glass bead (Sigma) and 300 µL of PBS buffer were added to the tubes and the tubes were shaken at speed 4.0 m/s in a FastPrep 24™ for 20 s to release viable bacteria from the leaves. From this homogenate, 20 µL was directly plated on both *Sphingomonas* and *Pseudomonas* selective media to confirm viable bacteria were present at the end of the experiment (not shown). Next, 470 µL of DNA lysis buffer containing 3% SDS as described in “DNA extraction from bulk cultures and plant lysates” above was added to the remaining lysate to make a final SDS concentration of 1.88%. Garnet rocks (0.5 mL) were added to the lysate and DNA was extracted by bead-beating at speed 6.0 m/s in the FastPrep 24™, and purified as described earlier.

<u>hamPCR to determine bacterial load and composition the bacterial community:</u> hamPCR (*95*) was performed on seedlings from soil-grown plants using primers for the *A. thaliana GIGANTEA* gene as a host gene and primers for the V4 region of the 16S rDNA (Extended Data Table 1), using primers and cycling conditions recommended in (*95*). Metadata regarding sequenced hamPCR amplicons can be found in Extended Data Table 2 .

### Bacterial growth in whole vs. macerated leaves

<u>Sample preparation:</u> In spring 2021, ∼40 g each of wild *Arabidopsis thaliana* and *Brassica napus* leaves were collected from a local field site in Germany (community of Kusterdingen, 48°31’00.9"N 9°06’34.9"E) with sterile scissors and tweezers, and kept them cool on ice. The larger *B. napus* leaves were trimmed into pieces no larger than ∼5 cm^2^. Upon returning to lab, the *A. thaliana* and *Brassica napus* leaves were washed in separate batches with copious amounts of distilled water and finally autoclaved sterile water to remove as many dirt and sediment-associated microbes as possible. From each species, 9 large aliquots (∼2 g) and 24 small aliquots (∼0.5 g) were prepared. One of the 9 large aliquots was macerated, diluted in an equal weight of PBS, and used for culturing bacteria on LB media. The other 8 large aliquots, to be used for repeated sampling of the macerated bacterial population, were each ground in a sterile mortar and pestle and each macerate was transferred to an empty petri plate. The 24 small aliquots were each placed with leaves adaxial (upper) sides up in a separate petri plate. All macerated leaves, whole leaves, and plated bacteria were incubated in 16° C in 16 hours of light.

For sampling on each of days 0, 1, and 2, eight of the small aliquots were sacrificed and ground in a mortar and pestle, and ∼0.3 g of each resulting homogenized sample was transferred to a pre-weighed screw-cap tube already containing 400 µL PBS and two 5 mm glass balls for weighing, CFU-counting, and 16S rDNA sequencing. Likewise, ∼0.3 g from the already macerated large lysate was removed using a sterile steel spoon and transferred to a pre-weighed screw-cap tube for the same procedures. At day 2 bacteria that had grown from the macerates that had been plated on LB plates were also collected.

<u>CFU counting:</u> Each screen cap tube containing 400 µL was ground for 20 seconds at speed 4.0 m/s in a FastPrep 24™ to release viable bacteria from the leaves. Then 400 µL of additional PBS was added and 20 µL was plated in a dilution series on LB media with 100 µg/mL cycloheximide and 100 µg/mL nitrofurantoin to select *Pseudomonas*.

<u>16S rDNA sequencing:</u> After the fresh lysate had been removed for CFU counting, 65 µL of 20% SDS (∼1.625 % final) and 0.5 mL garnet rocks were added. This was processed as described in the methods section “DNA extraction from bulk culture bacterial pellets and plant lysates”. The resulting DNA was amplified with V4 rDNA primers and sequenced as described in the methods section “16S rDNA V3-V4 amplicon sequencing”, with the exception that the forward primer was 515F as in (*10*), the annealing step was done at 50 °C, and the material was sequenced using paired 2×150 HiSeq 3000 reads. This sequencer was chosen because the experiment was small and it was expedient to spike the libraries into a lane of unrelated material. Because the 150 bp reads could not reliably be assembled into full V4 amplicons and a single 150 bp read alone could not provide sufficient resolution to distinguish SphASV1 and PseASV1 from other bacteria in the genus, the forward and reverse reads were simply concatenated and the concatenated sequences corresponding to PseASV1 and SphASV1 were identified and quantified separately. All other sequences were classified to the level of bacterial families based on the forward read alone. Metadata regarding these sequenced amplicons can be found in Extended Data Table 2 .

## Supporting information

Extended_Data_Table_1_PrimerSequences

Extended_Data_Table_2_SequenceMetadata

Extended_Data_Table_3_PlantSizeData

## Data availability

All data in this manuscript are deposited to the European Nucleotide Archive (ENA) under the project number PRJEB44136. At: https://www.ebi.ac.uk/ena/browser/view/PRJEB44136

The ENA accession numbers for individual raw reads and assemblies can be found in Extended Data Table 2 .

Custom scripts used are available at: https://github.com/derekLS1/ContrastingPatternsDominance

## Author Contributions

DSL planned the study. DSL, TLK, OS, RdP-J, and DW collected samples. DSL, RdP-J, PP, and KP processed plant samples, made sequencing libraries, and analyzed data. AB-G helped troubleshoot the agar plate assays. IB developed algorithms for plant phenotyping from photographs. WD helped with panX analysis. DSL, RdP-J, PP, and DW wrote the manuscript, with input from all authors.

## Acknowledgements

We thank Christa Lanz, Manuela Neumann, and Pablo Carbonell for assistance with Nanopore sequencing, Heike Budde for assistance with Illumina sequencing, and Haim Ashkenazy for assistance with the Nanopore genome assembly pipeline. Supported by Human Frontiers Science Program (HFSP) Long-Term Fellowships (LT000565/2015-L, D.S.L.; LT000348/2016-L, T.L.K.), ERC Advanced Grant IMMUNEMESIS (340602), the DFG through SPP Priority Program DECRyPT, and the Max Planck Society (DW).

