## Extended_Data_Figures for "Contrasting patterns of microbial dominance in the *Arabidopsis thaliana* phyllosphere"

### Extended Data Figure 1

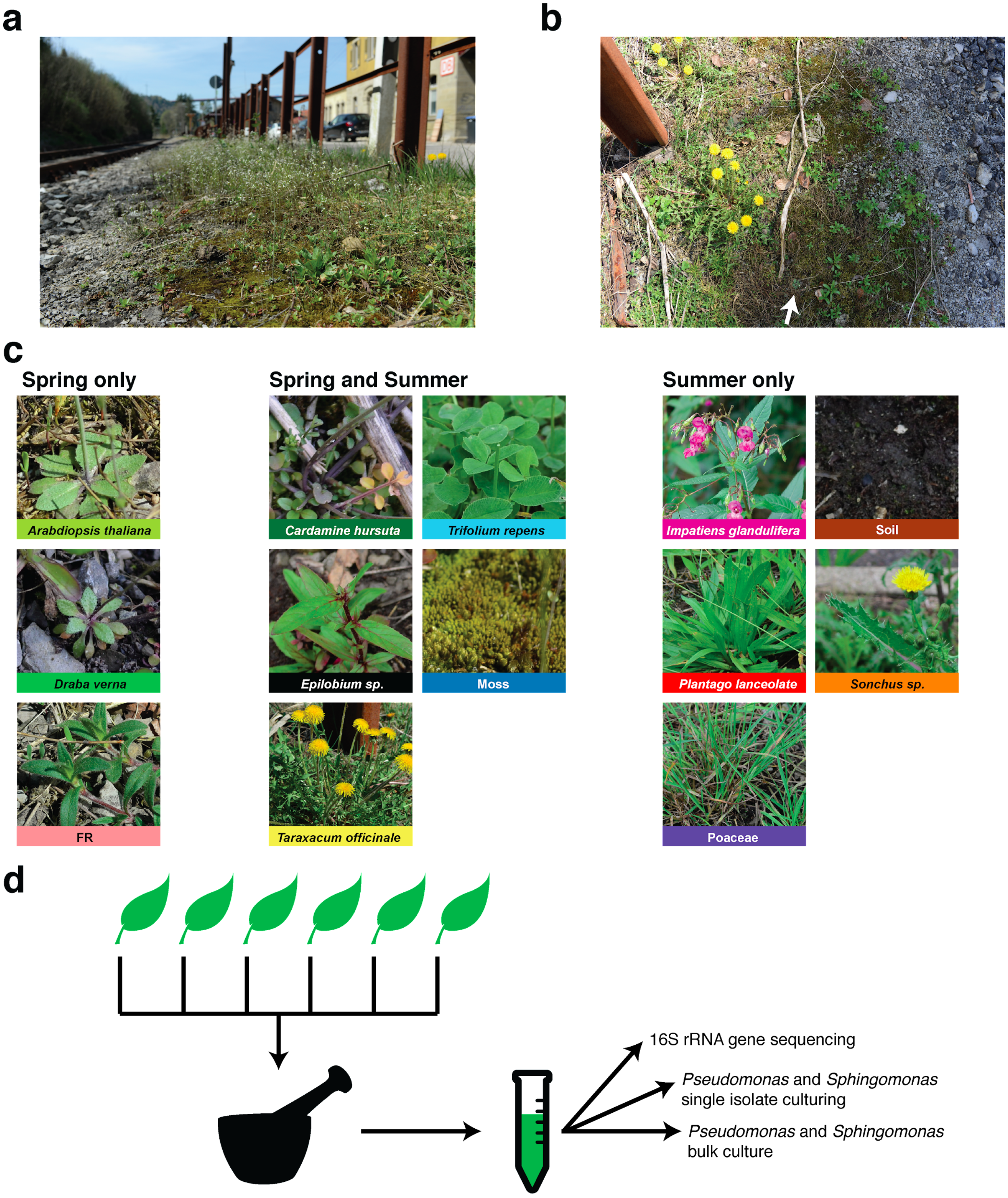


**Extended Data Fig. 1 | Sampling scheme. a**, The harvested area of Eyach, Germany on 20 April, 2018 (spring), viewed from the ground. The small white flowers are predominantly from *A. thaliana*. **b**, Part of the harvested area from **(a)** viewed from the air, showing *A. thaliana* (eg. arrow) and surrounding plants. **c**, Common plant species at the site were harvested only in the spring (left), both spring and summer (September 14, 2018, middle), and only summer (right). Soil was only harvested in the summer. When the plant species could be identified, the full Latin name is given. For all other plants, the genus or family is given. The species of moss remained unknown, as did the dicot plant named "FR", and these plants are referred to by their vulgar/code name. **d)** For each plant species, at least 1 leaf from at least 6 individuals per species was pooled to make a single sample. Seven such samples were produced per plant species. The leaves were ground in PBS in a mortar and pestle, and the macerate was mixed with glycerol to make a cryo-protected -80°C freezer stock to be used both for nucleic acid extraction and for culturing live bacteria.

### Extended Data Figure 2

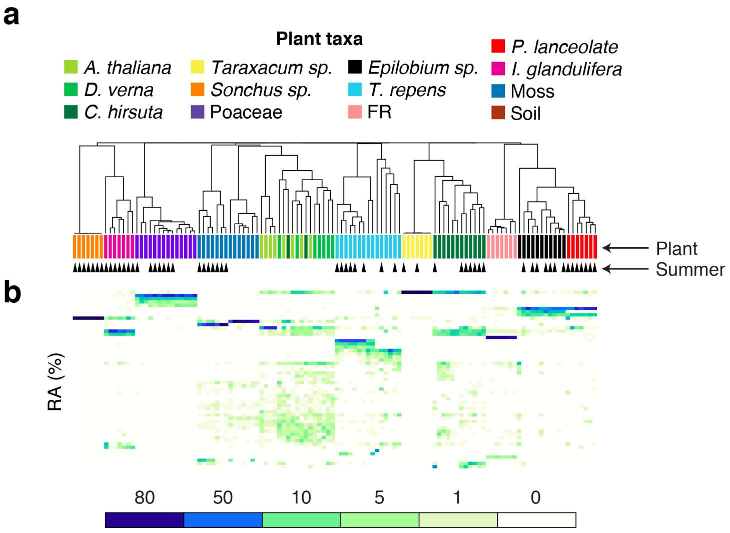


**Extended Data Fig. 2 | Analysis of sampled plant chloroplast sequences. a,** The relative abundances of 16S rDNA sequences classified as "chloroplast" were used to calculate pairwise Bray Curtis dissimilarity scores for all samples with ≥100 chloroplast sequences; the dissimilarity matrix was then used to cluster plants. **b,** Heatmap of chloroplast ASV relative abundances (RA%), showing the top 100 most abundant sequences. The tight clustering by plant species reveals that our identification of different plant taxa by their visual phenotype corresponded well with their genetics.

### Extended Data Figure 3

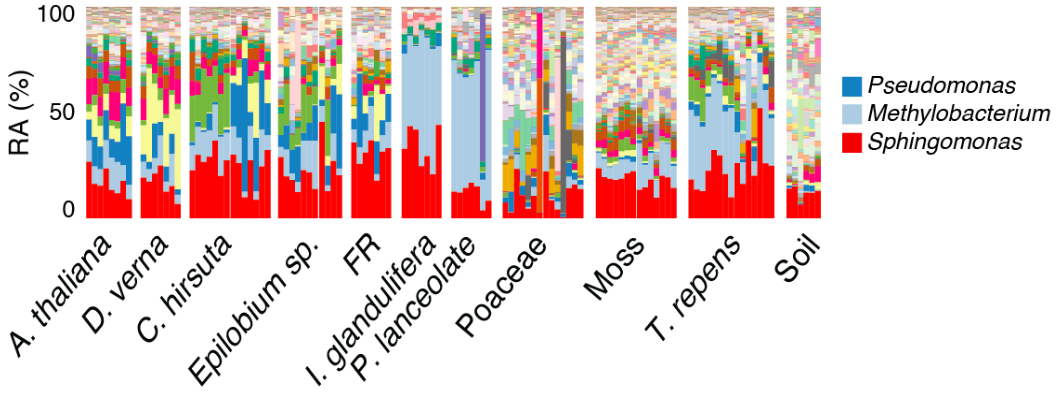


**Extended Data Fig. 3 | Relative abundance of major bacterial genera across plant species.** Stacked bars represent the relative abundances (RA%) of bacterial genera in the present study (in contrast to the bacterial families shown in Fig. 1) with the top 3 bacterial genera shown in reverse order in the legend at right. In contrast to Fig. 1, samples are grouped by plant species and not hierarchically clustered.

### Extended Data Figure 4

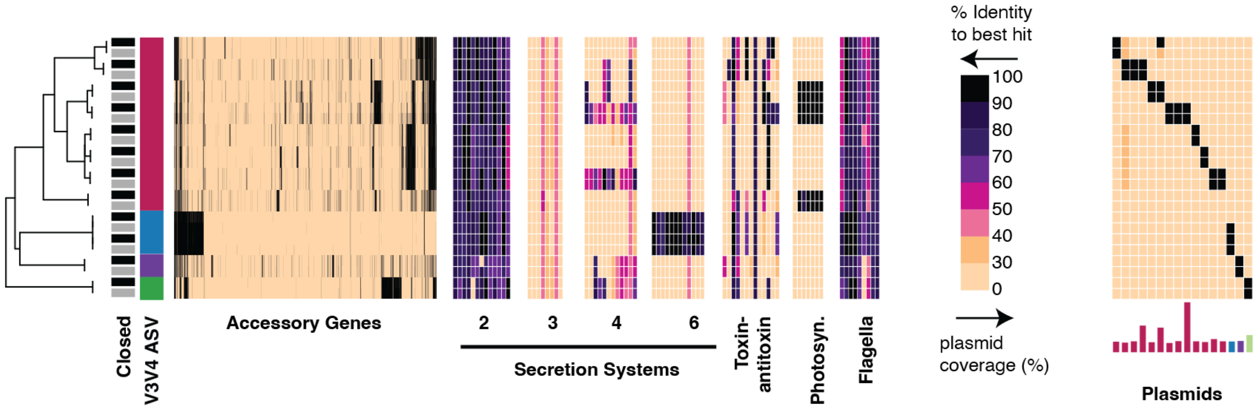


**Extended Data Fig. 4 | Closed genomes compared to their draft counterparts.** Nanopore-sequenced closed genomes (black boxes next to tree) are neighbors with their corresponding Hiseq draft genomes (grey boxes next to tree) in the core genome maximum-likelihood tree, and share the same gene presence and absence patterns with few exceptions. Most notably, one of the draft genomes has full coverage of a plasmid identified in other closed genomes, while the corresponding closed genome misses those genes, suggesting that perhaps in re-cultivation of the stock for Nanopore sequencing this plasmid was lost. The order of genomes (first Nanopore, then HiSeq) from top to bottom is: S216H113, S133H113, S127H113, S18H113, S213H113, S190H113, S230H113, S237H113, S380H113, S136H113, S132H113, and S337H113.

### Extended Data Figure 5

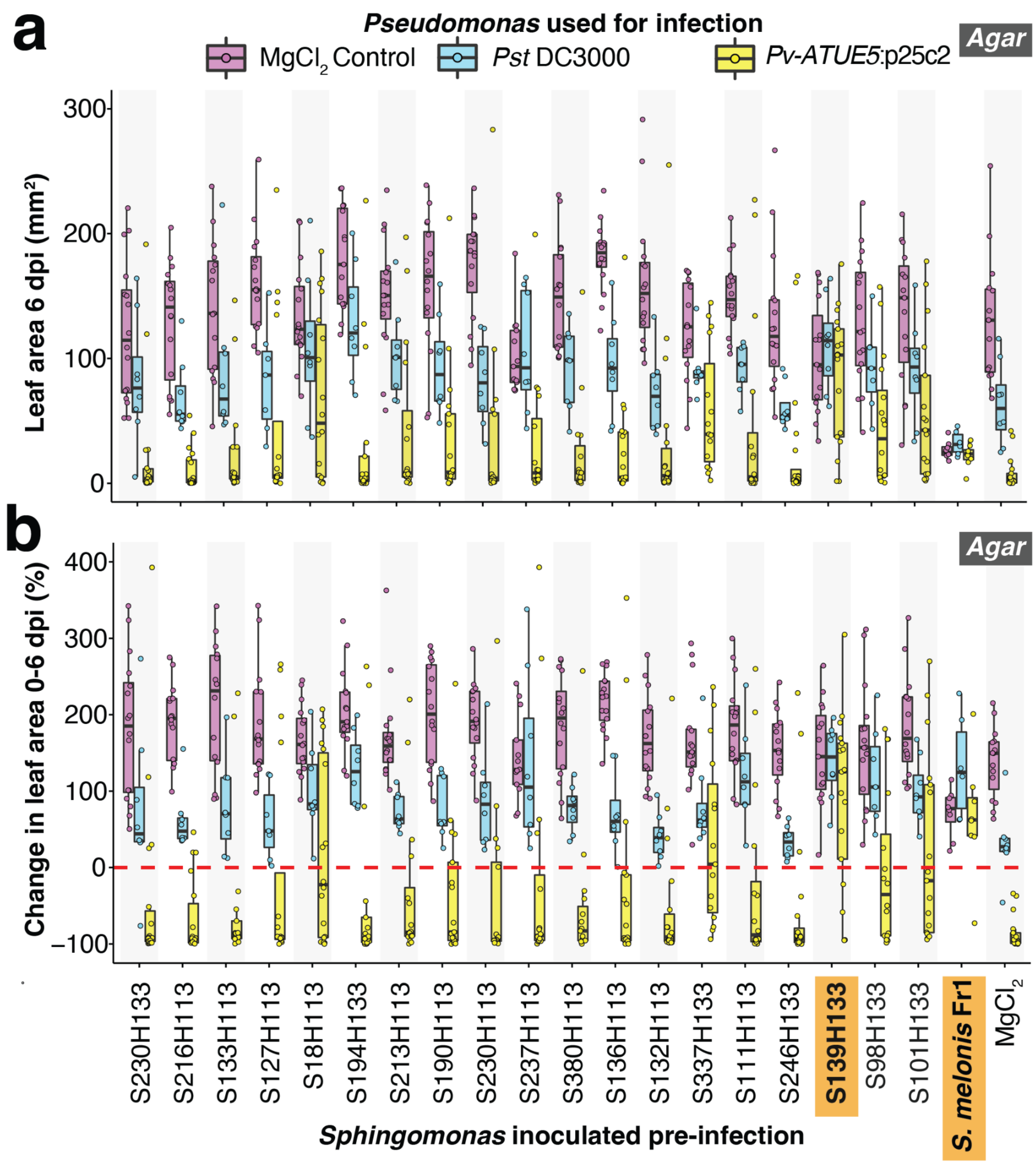


**Extended Data Fig. 5 | Gnotobiotic plant protection experiment for all tested strains. a**, In an independent experiment from that shown in Fig. 3a/3b, 19 local *Sphingomonas* isolates, *S. melonis* Fr1, or $MgCl_2$ buffer were used to pre-treat *A. thaliana* Ey15-2 seeds germinating in 24-well agar plates, and on day 10 seedlings were challenged with a $MgCl_2$ control, *Pto* DC3000, or *Pv-ATUE5*:p25c2 and monitored for 6 dpi. The Y-axis shows the rosette size at 6 dpi, with the dotted red horizontal line representing no change. **b**, Percent change in rosette size between 0-6 dpi. The orange highlighted *Sphingomonas* showed significant protection with no difference in symptoms from *Pst* DC3000 across this and the replicate shown in Fig. 3a/3b (FDR-adjusted Mann-Whitney U-test, $p > 0.05$).

### Extended Data Figure 6

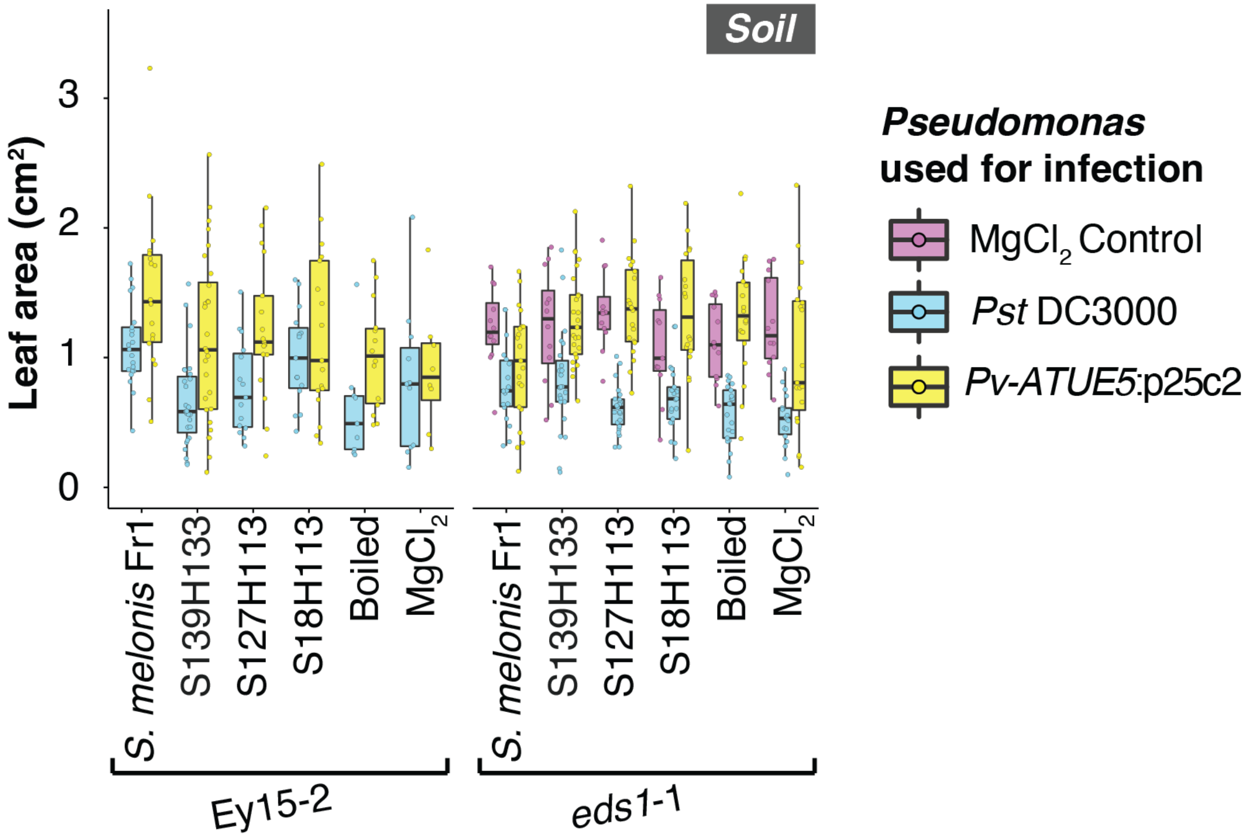


**Extended Data Fig. 6 | *Sphingomonas* protection against *Pst* DC3000 was not observed on soil.** Absolute sizes of rosettes of Ey15-2 and *eds1*-1 plants grown on soil for different *Sphingomonas* and *Pseudomonas* treatments, elaborated from Fig. 3e. The $MgCl_2$ control spray for *Pseudomonas* was omitted for the Ey15-2 genotype due to insufficient plants. Unlike on agar, *S. melonis* Fr1 treatment did not stunt the growth of seedlings. Plants not pre-treated with were larger than non-treated (buffer) plants for any genotype or *Pseudomonas treatment.*

### Extended Data Figure 7

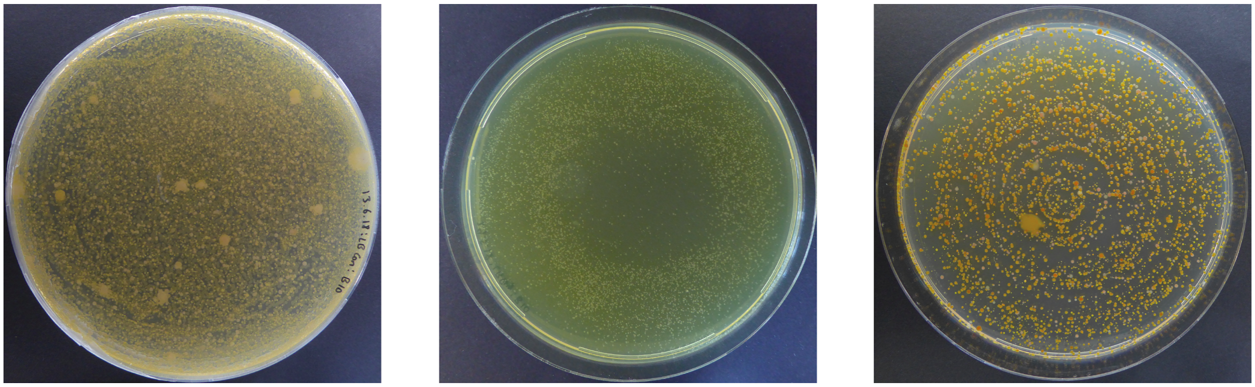

Control (LB + cycloheximide) | *Pseudomonas* selection (LB + cycloheximide + nitrofurantoin) | *Sphingomonas* selection (LB + cycloheximide + streptomycin)

**Extended Data Fig. 7 | Control and selective media plates for bulk culture of *Pseudomonas* and *Sphingomonas.*** From each lysate, 50 µL of glycerol stock corresponding to ~5 mg of original plant material was plated on selective *Pseudomonas* or *Sphingomonas* media. All three plates represent the same plant sample (*Draba verna*, HOST_PLANT_ID = ”Spring10”)

### Extended Data Figure 8

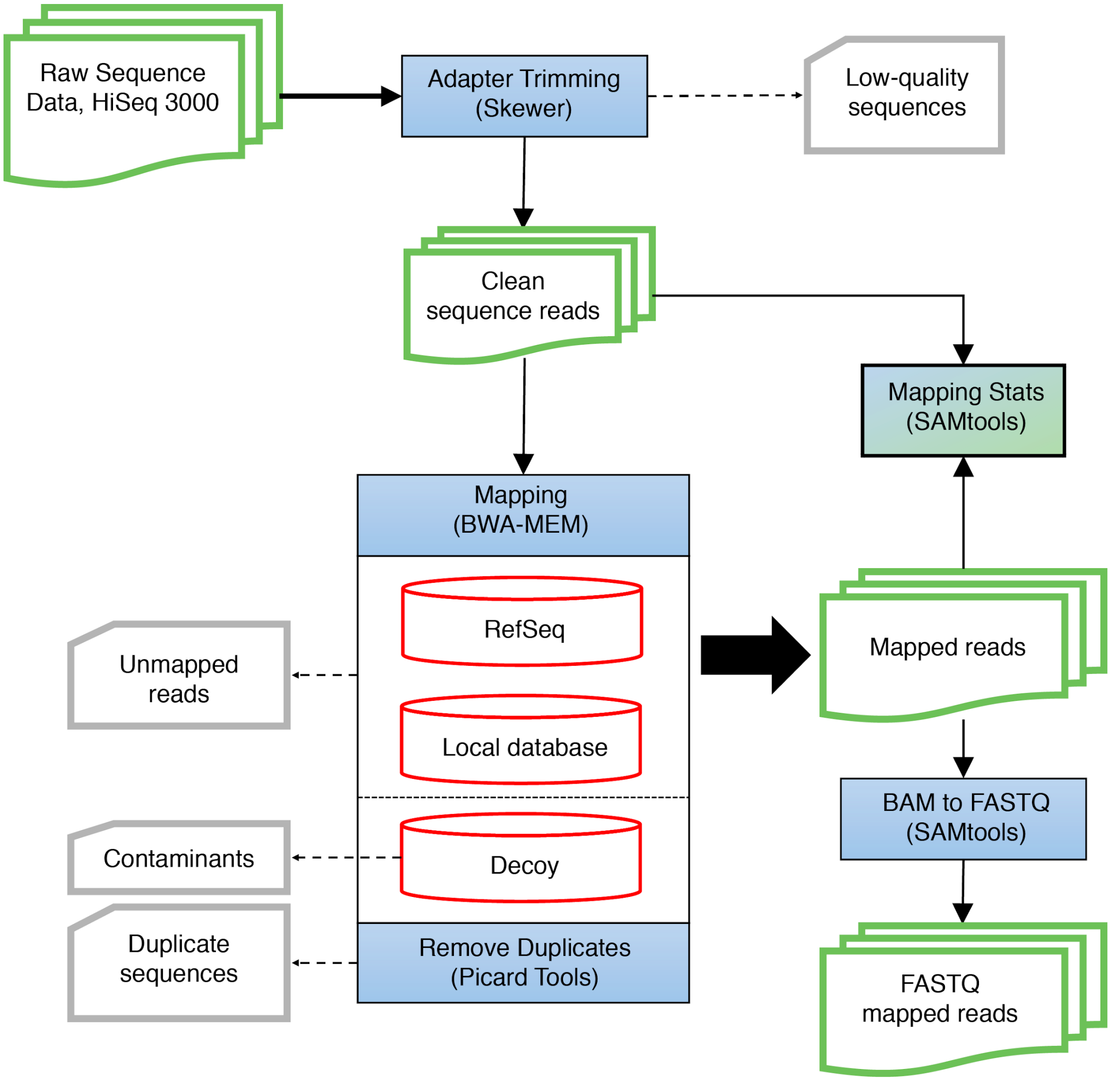


**Extended Data Fig. 8 | Metagenome analysis pipeline used in this study.** Raw sequences were trimmed, filtered, and mapped to a reference genome. The reads of interest were outputted as FASTQ files for further analysis.

### Extended Data Figure 9

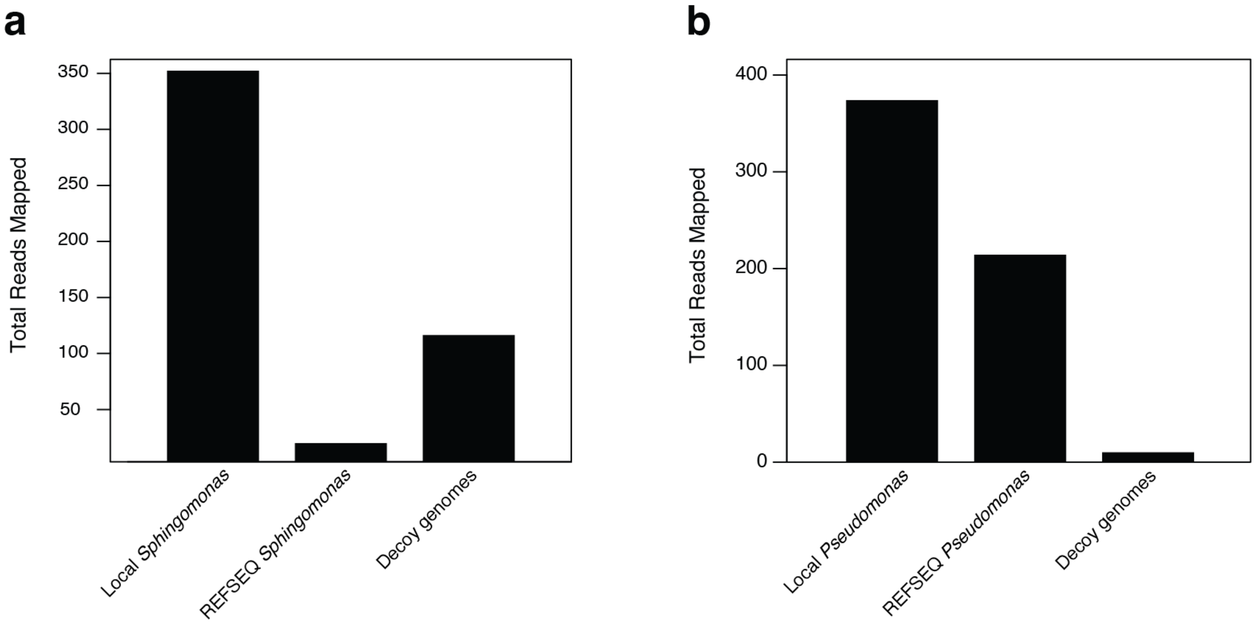


**Extended Data Fig. 9 | Metagenome reads mapping to local genomes, REFSEQ genomes, and DECOY genomes. a**, All bulk culture *Sphingomonas* metagenomes were mapped to local *Sphingomonas* genomes (left), Sphingomonas genomes from NCBI's REFSEQ (center) and decoy plant associated genomes from other genera to capture contaminants (right). Decoy reads represented 30% of all mapped reads. **b**, Same as **(a)**, but for *Pseudomonas*. Decoy reads represented 1.7% of all mapped reads.
